# Segmentation Matters: Recognizing the Cell Segmentation Challenge in Spatial Transcriptomics

**DOI:** 10.1101/2025.08.25.672145

**Authors:** Huasheng Yu, Anna Yao Carroll, Kevin Shen, Nathan Wu, Hanying Yan, Jingwei Xiong, Miguel Francisco Mercado, Eric A. Kaiser, Mayank Gautam, Mingyao Li, Wenqin Luo, Penn High Precision Pain Center

**Affiliations:** Department of Neuroscience, Perelman School of Medicine, University of Pennsylvania, Philadelphia, PA 19104, USA; Departments of Biostatistics, Epidemiology, and Informatics, Perelman School of Medicine, University of Pennsylvania, Philadelphia, PA 19104, USA; Graduate Group in Biostatistics, University of California Davis, Davis, CA 95616, USA; Department of Neurology, University of Pennsylvania, Philadelphia, PA 19104, USA

## Abstract

Probe-based *in situ* hybridization spatial transcriptomics has emerged as a state-of-the-art for neuroscience research. Accurate segmentation of neurons and non-neuronal cells, a critical step for downstream analysis, remains a big challenge. Using human sensory ganglia neurons as an example, we systematically explore this problem. We evaluated multiple automatic segmentation approaches using quantitative performance metrics and downstream results. We show that careful parameter tuning is essential for achieving accurate segmentation; however, even with optimized parameters, different models still yield distinct classes of errors. To mitigate these errors, we propose a manual quality check, which can validate and refine automated segmentation results. As a future direction, integrating multi-modal imaging data with tailored neural networks may further improve segmentation accuracy. In summary, our results argue that each automated segmentation method has distinct strengths, weaknesses, and characteristic error patterns; and that a manual review is necessary.

## Introduction

Spatial transcriptomics platforms based on *in situ* hybridization, such as commercial platforms including 10x Genomics Xenium, Vizgen MERSCOPE, NanoString CosMx SMI, and academic methods including MERFISH, seqFISH, and STARmap have emerged as transformative technologies for profiling hundreds to thousands of gene expressions in intact tissue sections at high spatial resolution^1–7^. These approaches are widely adopted, offering unprecedented insights into tissue architecture, cellular neighborhoods, and cell–cell interactions under physiological and pathological conditions. However, a key challenge in analyzing probe-based spatial transcriptomics data is the accurate segmentation of individual cells—especially specific target cell types—in densely packed, heterogeneous tissues.

Segmentation accuracy in spatial transcriptomics is not just a technical detail—it determines the reliability of downstream analyses and significantly affects biological interpretations. Manual cell segmentation is often considered the gold standard, but it is time- and labor-intensive and becomes impractical for large-scale or high-throughput studies. Several automated segmentation methods have been developed for various types of biological images^8–17^, and some of these approaches have been adapted for probe-based spatial transcriptomics.

Early MERFISH studies relied on nuclear staining (DAPI), often combined with polyA intensity basins to seed watershed or morphological filters for cell segmentation^18–20^. More recent work has shifted toward clustering-based algorithms (e.g., Baysor^13^) and deep learning approaches (e.g., Cellpose^12^). For Xenium, most published studies depend on the company platform, the Xenium Explorer, which also based on DAPI-seeded watershed. Despite the critical role of segmentation accuracy, most published studies applied these methods without systematic and independent validation of segmentation performance^3,11,19–25^. Given the diversity of current cell segmentation algorithmic strategies, from clustering approaches that operate on transcript coordinates to deep learning methods applied to pseudo-images, these methods are likely to differ in their strengths, limitations, and sources of error. This underscores the need for a systematic evaluation to guide the selection for segmentation method.

In this Viewpoint, we use human sensory ganglia neurons as a case to compare multiple segmentation methods, examining and comparing their performance, strengths, and weaknesses. We find that parameter tuning is critical for achieving optimal results with each method and present a framework how to systematically evaluate segmentation accuracy. We also propose to integrate a manual quality control step to reduce errors. As a future direction, integrating multi-modal imaging data with neural networks may further improve segmentation efficiency and accuracy. Our aim is to raise field-wide awareness of segmentation accuracy as a foundational step in spatial transcriptomics, to provide practical guidelines for selecting and refining segmentation approaches, and to offer perspectives for future advancements.

## Results

### Probe In-situ hybridization-based spatial transcriptomics

To evaluate segmentation strategies in spatial transcriptomics, we collected a set of 10x Genomics Xenium spatial transcriptomics datasets, including previously published human dorsal root ganglion sections (DRG1–DRG4)^26^ and a new human trigeminal ganglion (TG1) section (Figure S1A). Both DRG and TG datasets were profiled using a custom 100-gene probe panel containing marker genes for pan-sensory neurons, sensory neuron subtypes, satellite glial cells, and other non-neuronal cell types^26^. Sensory neurons were selected as the target cell type for segmentation.

Probe-based spatial transcriptomics workflows typically proceed through three main stages (Figure 1A): (1) spatial profiling of transcripts using technologies such as Xenium, (2) segmentation of cells or targeted cell types, and (3) downstream analyses, such as transcript quantification, clustering, and inference of cell-cell interactions.

**Figure 1.**
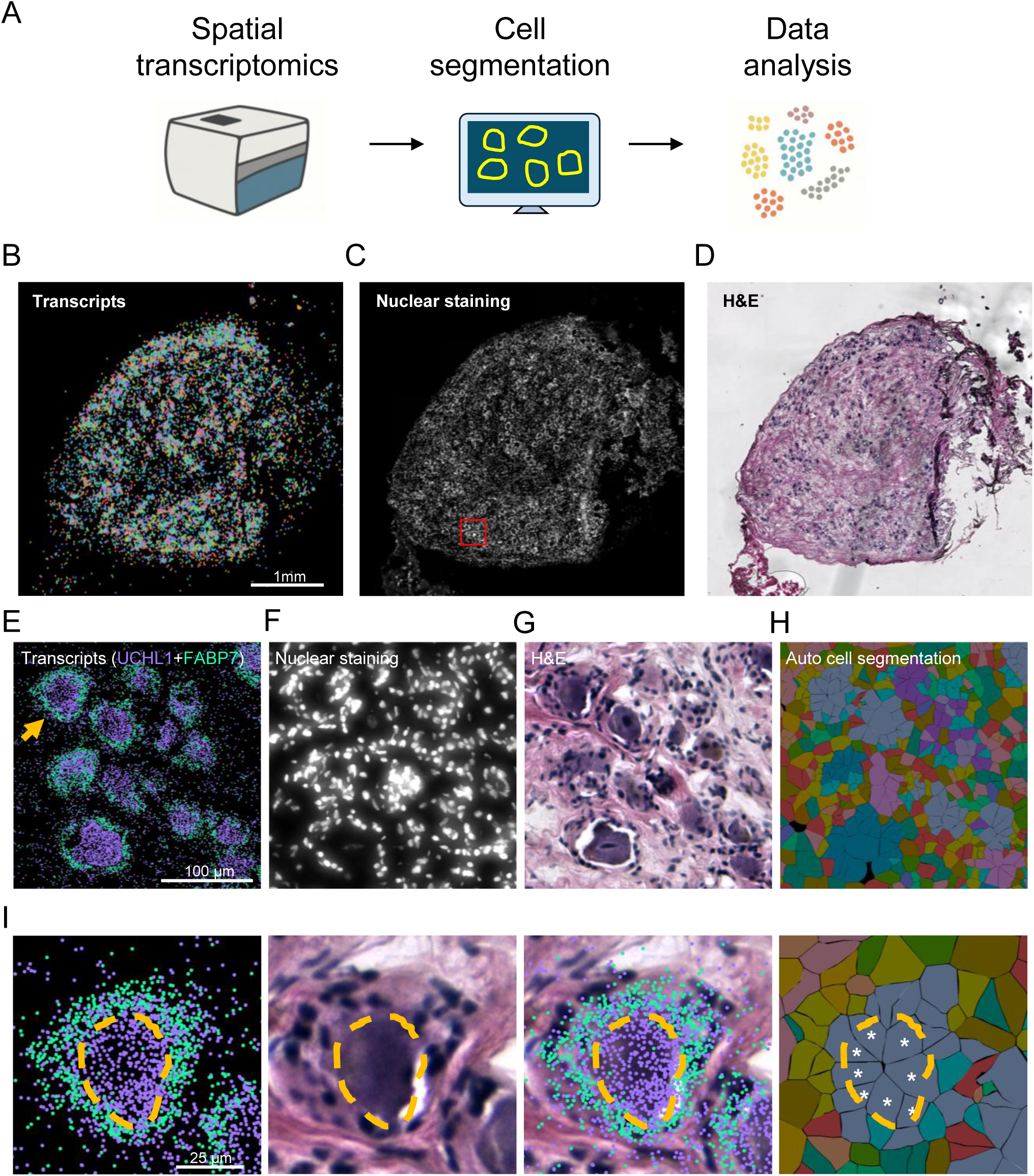
Probe-based *in situ* hybridization spatial transcriptomics. (**A**) Overall workflow of 10x Xenium spatial transcriptomics and data analysis. (**B-D**) An example of the 10x Xenium ST image (B), nucleus staining image (C) and H&E image (D) from human DRG. (**E-H**) A zoomed-in area showing the expression of transcript UCHL1 and FABP7 (E), nucleus staining (F), H&E staining (G), and Xenium automated cell segmentation (H). (**I**) A zoomed-in neuron from E-H.

Each Xenium dataset generated multiple image modalities, including pseudo-images derived from spatial transcript coordinates (ST images), nuclear staining (DAPI) images, and H&E-stained tissue images (Figure 1B–D). To delineate neuronal cell bodies, we used UCHL1—a pan-neuronal marker that forms dense transcript clusters—as the primary segmentation signal. The glial marker FABP7, which surrounds sensory neurons, was included to assist in boundary definition (Figure 1E).

We developed a Python-based preprocessing pipeline to convert transcript coordinates into ST pseudo-images. Each UCHL1 or FABP7 transcript was plotted as a single pixel on a black background, producing grayscale channels at 1 µm/pixel resolution. The corresponding H&E images were aligned to ST images using nuclear landmarks and resized to match dimensions (Figure S2A). Visual inspection of representative regions revealed tight alignment between UCHL1 transcript clusters and neuronal bodies in the H&E images (Figure 1E–G). Notably, sensory neurons exhibit much weaker DAPI nuclear staining compared to surrounding cell types (Figure 1F). Although Xenium platform provides an automated segmentation package, we observed frequent fragmentation of single neurons, partial boundaries, and over-segmentation (Figure 1H-I, S1A–B), underscoring the need for improved approaches.

### Manual segmentation and dataset preprocessing

Manual segmentation served as the ground truth and the foundation for both model evaluation and deep learning model training. Using ImageJ/FIJI, we manually annotated neuronal cell boundaries of all five sections. Most segmentation was guided by ST images, with the ROI Manager’s freehand tool used to delineate cell outlines. In cases where neurons were closely packed or had sparse transcript signals, we used H&E images to resolve boundary3 ambiguities and confirm the presence of cells (see Methods).

In total, we manually segmented 3,719 neurons across five tissue sections: 436 in DRG1, 375 in DRG2, 298 in DRG3, 306 in DRG4, and 2,304 in TG1. This annotated dataset provided the basis for model training and quantitative performance assessment.

For deep learning model development, we aligned ST and H&E images, removed peripheral regions, and cropped the images into 500 × 500-pixel patches. Corresponding manual cell annotation coordinates were adjusted accordingly. Cropped image patches from DRG1–3 were used to develop the models and were randomly split into training (80% of images), validation (10% of images), and internal testing (10% of images) (Figure S1A & S2A). The remaining two datasetsDRG4 and TG1were used only for testing to evaluate model generalization and compare different segmentation methods (Figure S2).

### Metrics used for model evaluation and comparison

To quantitatively compare segmentation performance, we used a set of complementary metrics (Figure S3). The Intersection over Union (IoU) is a widely used metric that measures the overlap between predicted and ground truth segmentations as the ratio of their intersection to their union. While IoU is informative, it does not penalize missed instances (false negatives) that have no matching prediction. To address this limitation, we also calculated precision, recall, and F1-score at an IoU threshold of 0.5 (referred to as F1-50). In this framework, a predicted mask with IoU > 0.5 relative to a manual (ground truth) mask is classified as a true positive; otherwise, it is considered a false prediction. These complementary metrics provide a more balanced view of model performance by incorporating both false positive and false negative.

To specifically assess boundary accuracy, we calculated F1-(50–95), which averages F1-scores across IoU thresholds from 0.5 to 0.95 in 0.05 increments. A model that achieves similar F1-50 but a higher F1-(50–95) score is better at delineating accurate cell boundaries. In addition, we computed pixel-wise precision, recall, and F1-score, which compare predicted and ground truth masks on a per-pixel basis (Figure S3A).

To better understand the nature of segmentation errors, we classified predictions into six categories: true positives (IoU > 0.5), partial errors (IoU < 0.5 with partial overlap), inner errors (IoU < 0.5 with predicted masks entirely within ground truth), excess errors (IoU < 0.5 with predicted masks larger than ground truth), false negatives (missed instances), and false positives (predictions with no matching ground truth) (Figure S3B).

The metrics described above were used for final model comparisons. However, during model training and parameter tuning, such as in developing YOLO and the Combo models, we used the model’s self-reported mean average precision at IoU 0.5 (mAP50) for internal performance monitoring and hyperparameter optimization.

It is also worth noting that different segmentation methods produce outputs in a variety of formats, including JSON files (Baysor), TXT files (DBSCAN, YOLO), CSV files (Cellpose), NumPy arrays (Combination model), and ROI files (ImageJ/FIJI). Therefore, we developed a pipeline to convert between these formats, enabling a standardized evaluation process (see Methods and GitHub link: https://github.com/taimeimiaole/Xenium-Segmentation).

### Different segmentation strategies

The primary data output of probe-based spatial transcriptomics is a set of spatial coordinates for each detected transcript molecule. To provide anatomical context, additional imaging data (such as nuclear stains like DAPI or histological H&E staining) are typically acquired alongside the transcript data. Broadly, two strategies can be employed for cell segmentation in spatial transcriptomics data: (1) clustering-based methods, which use the spatial distribution of marker gene transcripts to identify cells and approximate their boundaries, and (2) image-based methods, which apply deep learning models to segment cells in pseudo-images created from the transcript coordinates.

### Clustering-based methods

#### Baysor

Baysor is a probabilistic, Bayesian model-based tool for cell segmentation in spatial transcriptomics data. It assigns transcript molecules to cells based on spatial proximity and gene identity^13^. Optionally, Baysor can incorporate nuclear staining to help guide segmentation boundaries. However, in human DRG and TG sections, neuronal nuclear staining is much weaker than in non-neuronal cells and is barely visible (Figure 1F). Therefore, we excluded nuclear staining during Baysor segmentation optimization.

Several key parameters affect Baysor’s segmentation performance, particularly the choice of input transcripts, min_molecules_per_cell, and scale. The min_molecules_per_cell parameter defines the minimum number of transcripts required for a region to be considered a cell, while scale determines the expected spatial extent (i.e., the radius around the cell center within which transcripts are included). The Baysor segmentation workflow is illustrated in Figure S4A.

Since our goal was to segment sensory neurons, we focused on selecting pan-neuronal markers from our 100-gene panel. Based on visual inspection, *UCHL1* was selected for segmentation due to its better signal-to-noise ratio compared to other pan-neuronal markers, including *SYP*, *SLC17A6*, and *POU4F1*.

To assess the impact of min_molecules_per_cell and scale, we performed a grid search using different parameter combinations. First, with scale fixed at 15, we varied min_molecules_per_cell from 1 to 1000. The F1-50 score initially increased, peaked around 50– 100 molecules, and then declined (Figure S4B). Next, with min_molecules_per_cell fixed at 100, we varied scale from 1 to 100, observing optimal performance at a scale of approximately 10 (Figure S4C).

We applied the optimal parameter set for each DRG section and performed error classification analysis (Figure S4D). Across all four sections, the proportion of true positives remained consistently below 10%, indicating poor segmentation accuracy. The most common errors were partial segmentations, excess segmentations, and false positives, suggesting that Baysor struggled to distinguish background noise from true sensory neuron signals. We visually inspected Baysor’s results by overlaying predicted masks with manual annotations, which confirmed a high rate of overly large boundaries and false positive segmentations (Figure S4E). These findings suggest that Baysor may not be suitable for accurately segmenting human sensory neurons in this context.

#### DBSCAN

DBSCAN is another unsupervised clustering algorithm that can be applied to spatial transcriptomics data for cell segmentation^17^. To implement DBSCAN in our analysis, we followed the steps outlined in Figure 2A. DBSCAN groups transcripts based on local density. A point is classified as a core point if it has at least a specified number of neighboring points (MinPts) within a given radius (Eps). Points within this radius having fewer than MinPts neighbors are called border points, whereas points outside any core point’s neighborhood are considered noise points and are not assigned to any cluster during cell segmentation (Figure 2B).

**Figure 2.**
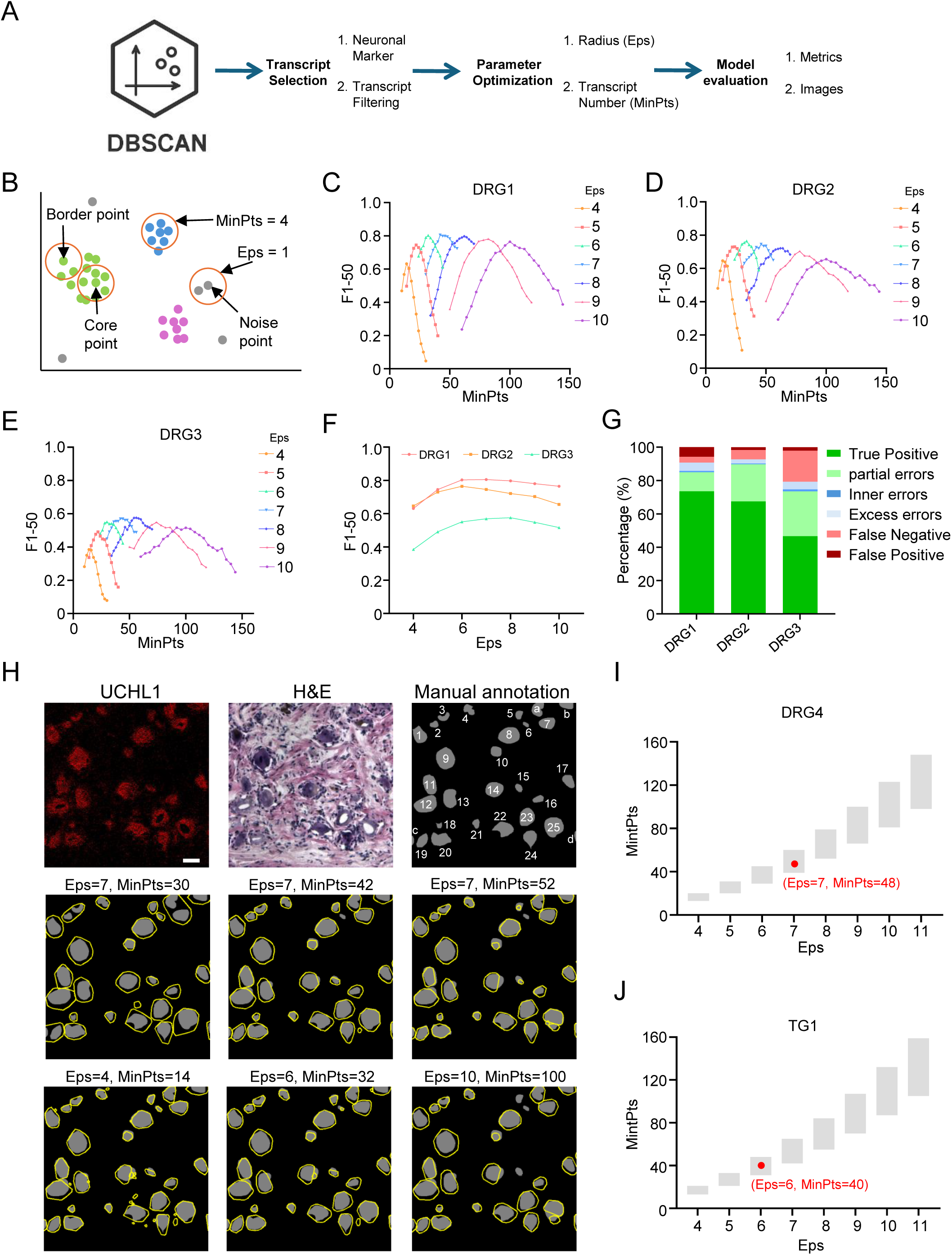
Unsupervised clustering-based segmentation by DBSCAN. (**A**) Overview of DBSCAN-based pipeline. (**B**) A cartoon illustrates key concepts of the DBSCAN algorithm. Core points are defined as points with at least a minimum number of neighbors (MinPts = 4) within a specified radius (Eps = 1, shown as orange circles). Border points fall within the neighborhood of a core point but do not themselves meet the MinPts requirement. Noise points (gray) are those that do not meet either condition. They are not assigned to any cluster. DBSCAN groups core and border points into clusters (shown in green, blue, and pink), while excluding noise points. (**C-E**) The effect of varying the DBSCAN parameters MinPts and Eps on segmentation performance, measured by F1-50 score, for DRG1 (C), DRG2 (D), and DRG3 (E). Each line represents a fixed Eps value, with F1-50 plotted across a range of MinPts values. (**F**) Maximum F1-50 scores achieved at each tested Eps value for DRG1–3 from (C-E), summarizing the best segmentation performance across a range of MinPts. (**G**) Bar plots summarizing the best segmentation results for DRG1-3. Segmentation results are grouped into six categories, including true positive, partial, inner, excess, false negative and false positive. (**H**) Representative DBSCAN segmentation results on a 500 × 500 µm region from DRG1 using different combinations of Eps and MinPts. Shown are the UCHL1 ST image, corresponding H&E image, manual annotation, and predicted segmentations. The optimal parameter combination (Eps = 7 and MinPts = 42 based on F1-50 score) is compared with five alternative combinations. Scale bar, 50 μm. (**I-J**) Predicted ranges of optimal Eps and MinPts values for DRG4 (I) and TG1 (J), based on our formula. The best-performing parameter combinations within the predicted ranges are indicated by red dots.

Similarly, we used pan-neuronal marker UCHL1 as the input for DBSCAN clustering. The two key parameters, Eps and MinPts, were systematically tested (Figures 2C–E). We observed a consistent trend across DRG1–3: for a fixed Eps, increasing MinPts initially improved F1-50 scores, peaking and then declining. Moreover, larger Eps values required larger optimal MinPts values, indicating that the two parameters are interdependent and should be jointly optimized. Among the conditions tested, Eps values between 6 and 8 generally yielded the best performance (Figure 2F). The segmentation error categories are summarized in Figure 2G.

To visualize the impact of these parameters on segmentation performance, we overlaid DBSCAN-generated segmentation outlines (yellow) with manual annotation (gray masks) (Figure 2H). Taking DRG1 as an example, the optimal parameter combination was Eps = 7 and MinPts = 42, achieving the highest F1-50 score of 0.8045 (Figure 2C). In a 500 × 500 µm zoomed-in example region of DRG1, manual annotation identified 25 neurons (excluding four boundary neurons from this analysis). The best Eps and MinPts combination correctly detected 24 neurons, missing only neuron 6 (false negative) and introducing one false positive cluster between neurons 24 and 25 (Figure 2H). When MinPts was reduced to 30 (Eps = 7), neuron 6 recovered. However, merging artifacts appeared, including merged clusters between neurons 2/3, which resulted in a reduced F1-50 score (0.5676). Increasing MinPts to 52 led to neuron 25 being fragmented into three smaller clusters and the boundaries of neurons 5, 10, 14, and 15 becoming smaller than real cell boundaries. Neuron 6 was missed again, and the F1-50 score dropped to 0.7600.

We also examined segmentation performance across a range of Eps values. At Eps = 4 and MinPts = 14 (optimal for Eps = 4), most neurons were detected, but the false positive rate was high, leading to an F1-50 of 0.6312. At Eps = 6 and MinPts = 32 (optimal for Eps = 6), performance was comparable to the optimal setting (F1-50 = 0.8030), although neuron 14 was split and neuron 6 was only partially detected. At Eps = 10 and MinPts = 100 (optimal for Eps = 10), cluster boundaries became a little bit larger than real cell boundaries, neurons 5, 6, and 15 were not detected and neuron 25 split, with the F1-50 declining to 0.7644. These results emphasize that DBSCAN performance is highly sensitive to the choice of Eps and MinPts. Thus, carefully tuning both parameters is critical for DBSCAN-based segmentation.

To guide parameter selection in real practice, we analyzed the segmentation results from DRG1– 3 and observed that optimal Eps and MinPts values are related to the cell size and transcript density (Figures S5A–C). We proposed a formula to estimate optimal Eps and MinPts range based on cell size (r_median_) and transcript density (d_median_):

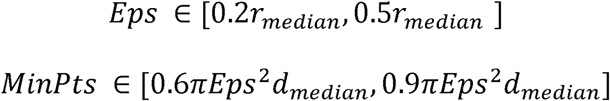

To determine the minimum number of neurons required for reliable estimation of r_median_ and d_median_, we performed 10,000 random subsampling iterations on 436 manually annotated neurons from DRG1, varying the subsample size from N = 5 to 60. For each iteration, we evaluated whether the known optimal combination (Eps = 7, MinPts = 42) fell within the predicted parameter range. As shown in Figure S5D, even with a sample size of 5 neurons, this optimal combination fell within the predicted range 83.80% of the time. This percentage increased rapidly with larger sample sizes: 96.05% at N = 10, 98.15% at N = 15, 99.47% at N = 20, and 99.96% at N = 30. These results suggest that robust parameter estimates can be obtained from a small subsample of annotated neurons.

To validate this estimation strategy, we applied it to two test datasets: DRG4 and TG1. For each, 20 neurons were randomly selected for manual annotation, and cell size and transcript density were estimated (Figures S5E–H). Using our proposed formula, the estimated Eps and MinPts ranges accurately captured the best Eps and MinPts combination: Eps = 7, MinPts = 48 for DRG4 (Figure 2I), and Eps = 6, MinPts = 40 for TG1 (Figure 2J). These results support the generalizability and practical utility of the parameter estimation for guiding DBSCAN segmentation in spatial transcriptomics data.

#### Deep learning-based methods

In addition to clustering-based segmentation methods, deep learning models are also widely applied in cell segmentation across various imaging modalities, including fluorescence microscopy, brightfield imaging, and electron microscopy (EM)^8–10,12,14–16^.

Although the raw output data for probe-based spatial transcriptomics only consists of transcript coordinates, they can be transformed into image-like representations suitable for image-based deep learning models. As previously described, UCHL1 and FABP7 were selected to generate training images. Details of training dataset preprocessing and preparation are described above (Figure S2A). To assess the feasibility of applying deep learning to cell segmentation in spatial transcriptomics, we evaluated three state-of-the-art models, including YOLO^27^, Cellpose^12^, and custom encoder-decoder architectures (Combo models).

#### YOLO

YOLO (You Only Look Once) is a user-friendly deep learning architecture capable of real-time object detection, pose estimation, classification, and instance segmentation^27^. YOLO is pretrained on large-scale datasets such as Microsoft COCO and ImageNet, making it highly transferable to new applications. To evaluate YOLO for ST image-based neuronal segmentation, we performed three key steps: model selection, training optimization, and model evaluation (Figure 3A).

**Figure 3.**
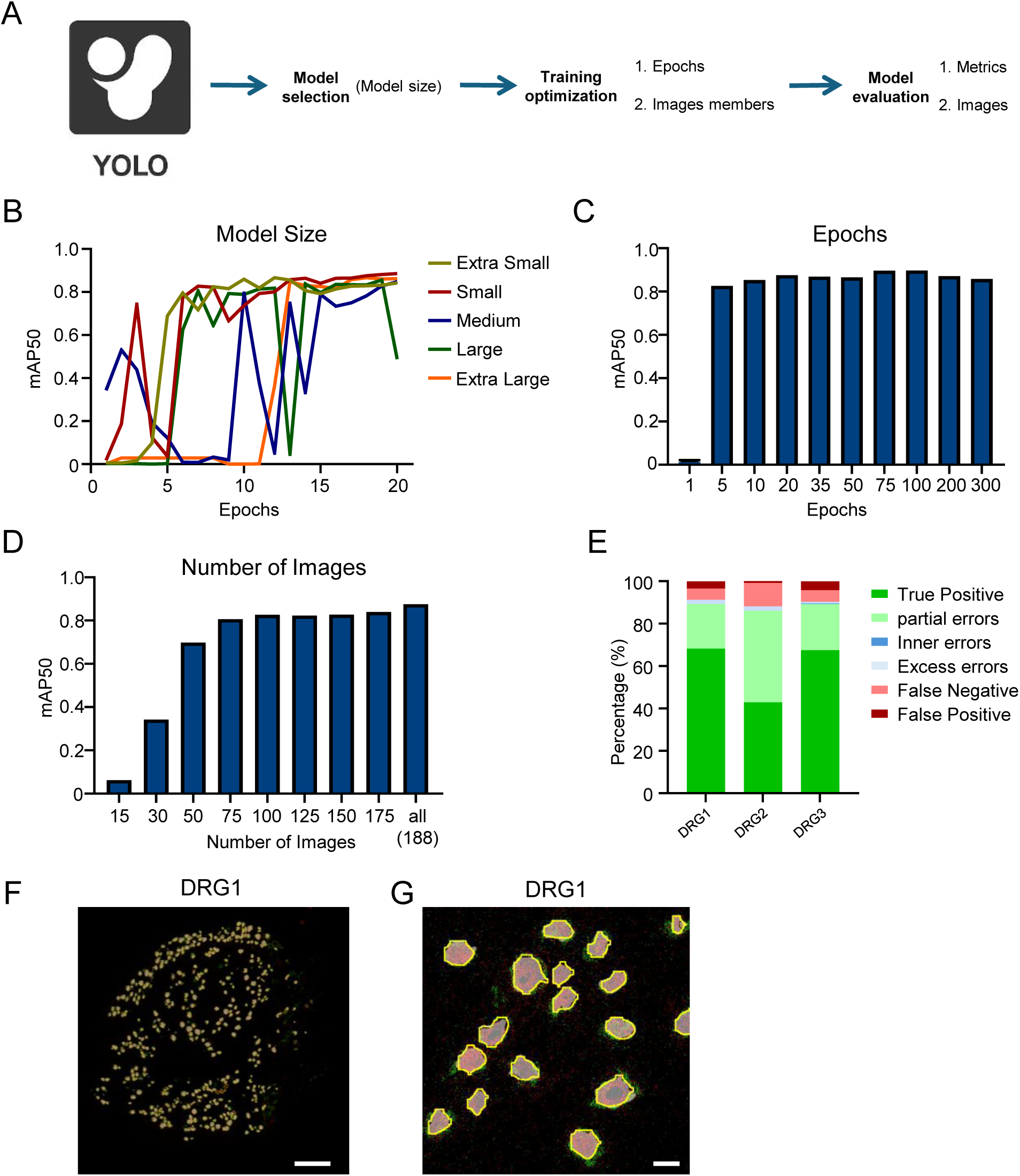
ST image-based segmentation by YOLO. (**A**) Overview of the YOLO segmentation workflow using ST images, including model selection, training optimization, and evaluation steps. (**B**) Effect of model size on segmentation performance. mAP50 values are shown for five YOLO model sizes (from extra small to extra large) trained on DRG1–3 ST images over 20 epochs. (**C**) Effect of the number of training epochs on model performance. The YOLO small model was trained on all 188 ST images with varying numbers of epochs. (**D**) Effect of training set size. The YOLO small model was trained for 20 epochs using subsets of an increasing number of images. (**E**) Segmentation error classification for the YOLO_ST model tested on DRG1–3 full tissue images. (**F–G**) Representative segmentation results for DRG1 whole image (F) and a zoomed-in area (G). Predicted neuron boundaries (yellow) overlaid on ST image. Scale bars: 500 µm in (F) and 50 µm in (G).

We first examined the impact of model size. YOLO (version 8) offers five segmentation model sizes: extra-small (3.4M parameters), small (11.8M), medium (27.3M), large (46.0M), and extra-large (71.8M). Larger models contain more parameters and can deal with more complex tasks, but they are also more computationally intensive. When trained on DRG1–3 for 20 epochs (DRG1-3 described in the dataset section), the small model demonstrated the best and most stable performance after 10 epochs (Figure 3B). Given the relatively small dataset and the focus on human sensory neurons, we selected the YOLO small model for all subsequent training steps due to both accurate and computational efficiency.

Next, we evaluated the effect of training epochs. Based on the default YOLO training epoch amount (100), we tested a range from 1 to 300 epochs to identify the optimal number for our dataset (Figure 3C). Performance, measured by internal parameter mAP50, increased rapidly and plateaued after approximately 5 epochs. The model achieved peak performance around 100 epochs, with no further gains and slight performance drops after 100 epochs. Based on this result, we used 100 epochs for the final training and test.

To determine whether our dataset was sufficiently large to train a robust model, we assessed the effect of training image quantity on performance. DRG1–3 contained 188 cropped training images. We trained the model using subsets of increasing size (15, 30, 50, 75, 100, 125, 150, 175, and 188 images) while keeping the test and validation sets fixed for consistent comparison (Figure 3D). Each larger subset included all images from the smaller subsets. As expected, performance improved with more training data, reaching a plateau after approximately 75 images. Training with all 188 images yielded the highest mAP50 score of 0.875.

Using these optimized parameters—small model size, 100 training epochs, and all 188 images— we trained the final YOLO segmentation model (YOLO_ST) and evaluated its performance on full tissue sections of DRG1–3. As shown in Figure 3E, YOLO achieved approximately 90% detection accuracy across all three DRG sections. However, a proportion of the predictions had less accurate boundaries, with partial overlaps with ground truth (IoU < 0.5). DRG2 showed the highest rate of partial errors, reflecting inter-sample variability. False negatives (missed cells) plus false positives (wrong predictions) were around 10% for all three DRG sections. A representative full-section prediction for DRG1 and a zoomed-in region are shown in Figures 3F and 3G, where the predicted neuron boundaries (yellow) are overlaid on the original ST image alongside the manual annotations (gray masks).

#### Cellpose

Cellpose is a widely used and well-established model specifically designed for cell segmentation^12^. Unlike YOLO, Cellpose incorporates image preprocessing prior to neural network training and inference. Specifically, it applies a simple diffusion algorithm to the ground truth masks, generating spatial gradients in the horizontal and vertical directions. Additionally, it produces a cell probability map that quantifies the likelihood of each pixel belonging to a cell. To evaluate Cellpose in the context of spatial transcriptomics, we applied a similar workflow to YOLO, fine-tuning the training and inference parameters (Figure 4A).

**Figure 4.**
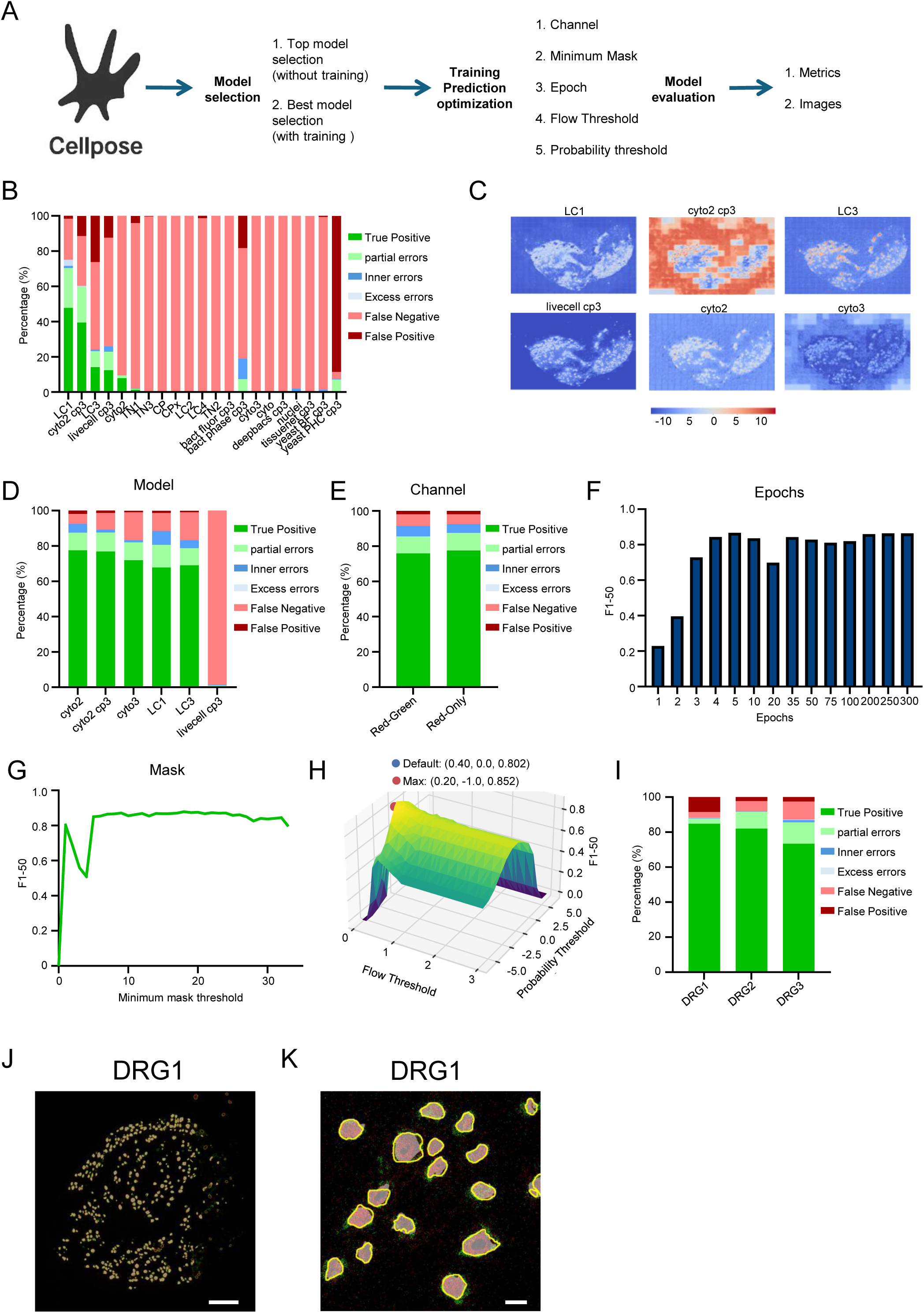
ST image-based segmentation by Cellpose. (**A**) Overview of the Cellpose segmentation workflow. The process is divided into three main stages: (1) initial pretrained model selection, (2) training and prediction parameter optimization, and (3) final model evaluation using both metrics and visual inspection. (**B**) Segmentation performance evaluation of all initial pretrained models based on prediction categories applied to DRG4 without training. (**C**) Visualization of segmentation probability maps from initial top pretrained models (LC1, cyto2 cp3, LC3, livecell, cyto2, and cyto3) applied to DRG4. (**D**) Segmentation performance on DRG4 of the six top pretrained models after training with DRG1-3 dataset. (**E**) Comparison of segmentation results trained with both red (UCHL1) and green (FABP7) channels (RG) versus red only (UCHL1) channel (R) using cyto2 model. (**F**) Effect of training epochs on segmentation performance for the cyto2_R model trained with varying epoch numbers. (**G**) Segmentation performance trained for 300 epochs on cyto2_R model across a range of minimum mask thresholds. (**H**) 3D plot of F1-50 scores for cyto2_R_18_mask model predictions across a range of flow and probability thresholds, illustrating the optimal parameter combination (Max) compared to the default. (**I**) Segmentation error classification for the Cellpose_ST model applied to DRG1–3 full tissue images. (**J-K**) Representative segmentation results for DRG1 whole image (J) and a zoomed-in area (K). Predicted neuron boundaries (yellow) overlaid on ST image. Scale bars: 500 µm in (J) and 50 µm in (K).

Cellpose offers a collection of pretrained models developed across diverse imaging contexts. To identify the most suitable model for our ST images, we first evaluated all available pretrained models on the DRG dataset without additional training. As shown in Figure 4B, performance varied substantially across these pre-trained models. Visualization of the raw cell probability maps illustrated how each model differentiated cells from the background (Figure 4C). Based on this initial screening, we selected six top-performing candidates—LC1, cyto2_cp3, LC3, livecell_cp3, cyto2, and the default cyto3—for further training on our DRG1–3 dataset. Among pretrained models, LC1 yielded the highest performance (Figure 4B). However, after adapting to our dataset through additional training (Figure 4D), cyto2 outperformed the others and was thus selected for the final training and testing.

Cellpose supports both single- and dual-channel segmentation modes, where each channel represents a different fluorescence signal. The second channel is an optional nuclear channel originally used for segmentation of the nucleus. In our ST pseudo-image, the red channel represents to the neuronal marker UCHL1, and the green channel corresponds to the glial marker FABP7. In single-channel mode, only the most informative red channel was used. In dual-channel mode, both red and green channels were used. Using the cyto2 model and default training parameters, segmentation based on the red channel alone (UCHL1 signals) slightly outperformed the dual-channel configuration (Figure 4E).

We also tested training performance across a wide range of epochs (Figure 4F). Model performance improved rapidly in the early epochs and then plateaued. Training for 5 epochs yielded the highest F1-50 score (0.8668), while 250 epochs achieved a comparable F1-50 (0.8642) but a slightly higher F1-(50–95) score (0.5401 vs. 0.5093) (Figure 4F, Supplementary Table 1), indicating improved boundary accuracy. Based on this result, we selected 250 epochs for the final training.

Cellpose includes a min_masks parameter that defines the minimum number of annotated cells required in an image to be included for training. We tested min_masks thresholds ranging from 1 to 33. Interestingly, performance varied considerably at low thresholds (1–3), stabilized after 4 and 26, and slightly declined at higher values (Figure 4G). The best-performing threshold was 18 masks, highlighting the importance of tuning this parameter for optimal performance.

For inference, Cellpose allows tuning of two critical parameters: flow threshold and cell probability threshold. A combinatorial test of these parameters revealed that optimized values improved the F1-50 score (0.852), compared to the default setting (flow threshold = 0.4, cell probability threshold = 0) (Figure 4H).

By combining all optimized parameters, we trained the final Cellpose model (Cellpose_ST) and tested it on the three full DRG datasets (DRG1–3). As shown in Figure 4I, the trained model achieved high segmentation accuracy, with most predictions classified as true positives and only a small proportion identified as partial errors, false negatives, or false positives. For the visualization, Cellpose predictions on DRG1 were overlaid with manual annotation onto full-section (Figure 4J) and a zoomed-in region (Figure 4K).

#### Combination model

In addition to YOLO and Cellpose, we explored other deep learning neural networks. For this purpose, we systematically evaluated the open-source Segmentation Models library (Version 0.3.4)^28^, which offers a broad collection of architectures and encoders. The architectures tested include nine widely used designs—Unet, Unet++, MAnet, Linknet, FPN, PSPNet, PAN, DeepLabV3, and DeepLabV3+—and ten families of encoders ResNeSt, ResNeXt, ResNest, Res2Net, RegNet, SE-Net, DenseNet, Inception, EfficientNet, and VGG. These architectures vary in complexity and design principles, and the encoders span lightweight to high-capacity designs, offering flexible combinations suited for various image characteristics. Compared to YOLO (anchor-based detection) and Cellpose (vector field-based mask regression), these models perform pixel-wise segmentation and were thus worth exploring for the ST images.

To systematically test this library collection and select the best architecture and encoder for our case, we used the same DRG1–3 ST images for training across all combinations. Specifically, we paired each of the nine architectures with each available encoder (72 combinations total) and evaluated each combination’s performance by averaging its last five self-reported validation mAP50 metrics. Among the architectures, Unet, MAnet, Linknet, and Unet++ achieved the highest average mAP50 scores (Figure 5A). Similarly, when comparing encoder families across all architectures, ResNeXt, Res2Net, and VGG were the top-performers (Figure 5B). Due to compatibility issues between Unet++ and most encoders, Unet++ was excluded from subsequent tests.

**Figure 5.**
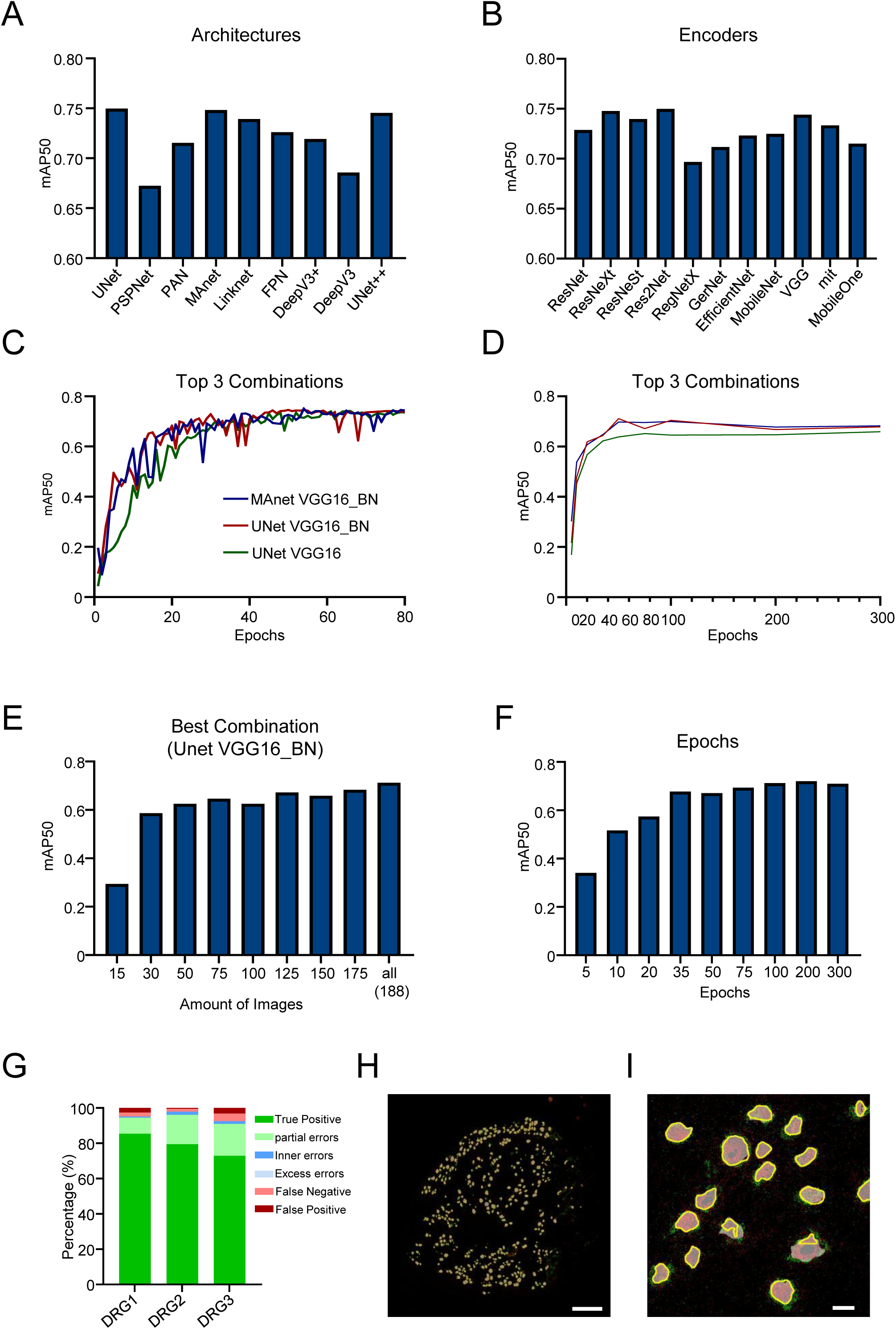
ST image-based segmentation by Combo model. (**A**) Average mAP50 scores across various model architectures (e.g., UNet, PSPNet, PAN, etc.) averaged over all encoder combinations. (**B**) Average mAP50 scores across different encoders (e.g., ResNet, VGG, MobileNet) averaged over all architectures. (**C**) Validation mAP50 growth curves over 80 epochs for the top three performing architecture-encoder combinations: MAnet + VGG16_BN, UNet + VGG16_BN, and UNet + VGG16. (**D**) Growth of test mAP50 values for each of the top 3 combinations when trained on various epochs (5, 10, 35, 50, 75, 100, 200, and 300). (**E**) mAP50 scores of the best-performing combination model (UNet + VGG16_BN) when trained with different numbers of training images. (**F**) Performance of the best-performing combination model (UNet + VGG16_BN) when trained for varying numbers of epochs. (**G**) Segmentation error classification for the Combo_ST model when tested on DRG1–3 full tissue images. (**H-I**) Representative segmentation results from the Combo_ST model for DRG1 whole image (H) and a zoomed-in area (I). Predicted neuron boundaries (yellow) overlaid on ST image. Scale bars: 500 µm in (H) and 50 µm in (I).

Next, we trained all combinations between the top three architectures and the top three encoder families (at all available model sizes). We averaged the validation mAP50 over the final five epochs for each combination. The combinations Unet + VGG16_BN, Unet + VGG19, and MAnet + VGG16_BN achieved the top 3 scores. Figure 5C displays the growth in validation mAP50 of the top three combinations over 80 epochs. Each of these three combinations were then trained on various epoch amounts to determine the optimized epoch number and best performing model. Figure 5D displays the trend in test mAP50 values for each of the top 3 combinations when trained on different epoch values. While Manet + VGG16_bn achieved two of the highest mAP50 scores, its performance seemed inconsistent. To illustrate, this combination achieved a 0.71 and 0.70 mAP50 score when trained on 50 and 100 epochs, respectively. Yet, leading up to the 0.71 metric, there was an increase in mAP50 of 0.07 and then after 0.71, there was a decrease of 0.04. Overall, the performance of MAnet + VGG16_bn seemed to fluctuate too often. Opposingly, Unet + VGG16_bn showed a continuous increase in mAP50, peaking at 100 epochs with an mAP50 score of 0.699. Since this was only a 0.01 difference in mAP50 compared to Manet+VGG16_bn’s highest metric and the combination showed greater consistency, we chose Unet + VGG16_bn as our final, top performing combination. The third combination of Unet + VGG19 performed worst, achieving only a 0.658 as its highest accuracy score.

The final model (Combo_ST) was tested on the whole DRG1–3 images. It achieved high accuracy, with most predictions being true positive and only a small fraction classified as partial errors. False negatives and false positives are minimal (Figure 5G). Overlaying the predicted masks on the original DRG1 ST image along with manual annotation demonstrated good agreement between the predicted and actual cell boundaries (Figures 5H–I).

### Leveraging multimodal imaging for improved segmentation

While most segmentation efforts in spatial transcriptomics rely on a single image modality, combining complementary sources of information can, in principle, improve model accuracy. Multimodality approaches—where multiple imaging channels or modalities are jointly used for training and prediction—remain underexplored in probe-based spatial transcriptomics. With the increasing availability of high-quality multimodality datasets, there is now an opportunity to test whether deep learning architecture can effectively integrate multi-modal inputs for better segmentation performance. Here, we hypothesize that combining ST images with H&E images would facilitate model accuracy, due to additional information on boundaries and cell presence.

#### H&E images

As mentioned before, in addition to ST images, we also acquired H&E images following Xenium spatial transcriptomics. To first evaluate whether H&E images alone could achieve accurate segmentation using deep learning models, we trained and evaluated both YOLO and Cellpose using H&E images, following the same pipeline for parameter optimization.

For YOLO, we systematically tested different model sizes, training epochs, and numbers of training images (Figure S6A–D). The small YOLO model achieved the highest accuracy (Figure S6B). Model performance improved with more training epochs and plateaued around 100 epochs (Figure S6C). Increasing the number of training images also led to higher accuracy, with the best performance obtained when using all 188 available images (Figure S6D). The final YOLO_H&E model was trained using the optimal parameters identified through this process. Error analysis showed that most predictions were true positives, though some partial and false negative predictions remained (Figure S5E). Representative segmentation examples are shown in Figure S6F–G.

Similarly, Cellpose models were optimized across multiple parameters, including different pretrained models, input channels, training epochs, and segmentation thresholds (Figure S6A–G). Among the pretrained models, CPx yielded the best performance (Figure S6A-C). Using the red single channel is slightly better than using the red-green double channels (Figure S6D). Performance increased with more training epochs, reaching a plateau around 100–200 epochs (Figure S6E). The minimum mask threshold had a substantial impact on performance, with the F1-50 score stabilizing when the threshold was larger than 2 (Figure S6F). Additionally, prediction parameters such as flow threshold and probability threshold were systematically tuned, with optimal settings showing in the 3D plot compared to default value (Figure S6G). Error analysis and representative segmentation results are shown in Figure S6H–J. In summary, while both YOLO and Cellpose could segment cells from H&E images, their performance was slightly inferior to models trained on ST images (Figure 3-4).

#### Combine both ST images and H&E images for the model training

ST images and H&E images provide complementary information for delineating cell boundaries. ST images, using neuronal marker UCHL1 and glia cell marker FABP7, help localize neuronal cell bodies and delineate the cell boundaries, while H&E images offer additional histological information that is especially useful when cells are densely packed or difficult to distinguish in ST images. Notably, H&E images were frequently referenced during manual annotation to determine the boundaries of closely attached cells. Therefore, we tested whether combining ST and H&E images would improve model performance compared to using ST images alone.

YOLO models only accept a single RGB image as input, so we generated merged images by integrating information from both modalities. As illustrated in Figure S8A, we first inverted the H&E image to transform the background from white to black, preventing interference with other channels. Then we split the H&E image into red, green, and blue channels. We selected the red channel for subsequent use due to it visually having the best contrast for neuronal boundaries. UCHL1 and FABP7 were already single-channel grayscale images, which we directly assigned to the blue and green channels, respectively. These three channels were then merged to generate the final composite RGB image.

Following the YOLO training pipeline, we systematically optimized model size, training epochs, and the number of training images (Figure S8B–D). We found that the small YOLO model trained for 100 epochs using the complete image set (188 images) yielded the best performance. Using these parameters, we compared models trained on ST image-only, H&E image-only, and merged images. As shown in Figure S8E, the YOLO model trained on merged images (YOLO_Merged) consistently achieved higher mAP50 scores throughout training, outperforming both ST-only and H&E-only models. Evaluation on DRG1–3 showed that YOLO_Merged achieved a higher mAP50 (Figure S8E) and higher average true positive rate compared to the YOLO_ST model alone (Figure S8F). The representative segmentation results are shown in Figure S8G–H. These results suggest that merging ST and H&E images can benefit YOLO-based segmentation.

In parallel, we applied the same merged image strategy to Cellpose models. Using the same merged images (one channel for H&E, one for UCHL1, and one for FABP7), we tested a wide range of training parameters, including different pretrained models, channel assignments, number of epochs, and minimum mask thresholds (Figure S9A–E). We also tuned prediction parameters such as flow and probability thresholds (Figure S9F). Despite these efforts, the performance of Cellpose on the merged dataset was notably poor, significantly worse than that of Cellpose trained on ST-only images (Figure S9G–J). This poor performance may be due to Cellpose being primarily optimized for high-contrast fluorescence images, making it less effective for segmenting cells in RGB-like composite images that include H&E.

### Evaluation of model performance on test datasets

To evaluate the generalization capability of different segmentation models, we tested them on an unseen DRG4 section from the same tissue type that was not included in the training set, and an unseen TG1, which originates from a different anatomical location (Figure S1A).

For DRG4, representative segmentation results and corresponding zoomed-in views are shown for four models: Cellpose_ST, YOLO_ST, Combo_ST, and DBSCAN (Figure 6A–D). All models correctly identified the most sensory neurons but differed in boundary accuracy. The Cellpose_ST and Comb_ST models produced smooth, rounded segmentation boundaries (Figure 6A&C) while YOLO_ST exhibited slightly boxy shapes (Figure 6B), and DBSCAN often generated slightly larger boundaries (Figure 6D). Similar trends were observed in the TG1 test dataset (Figure 6E–H).

**Figure 6.**
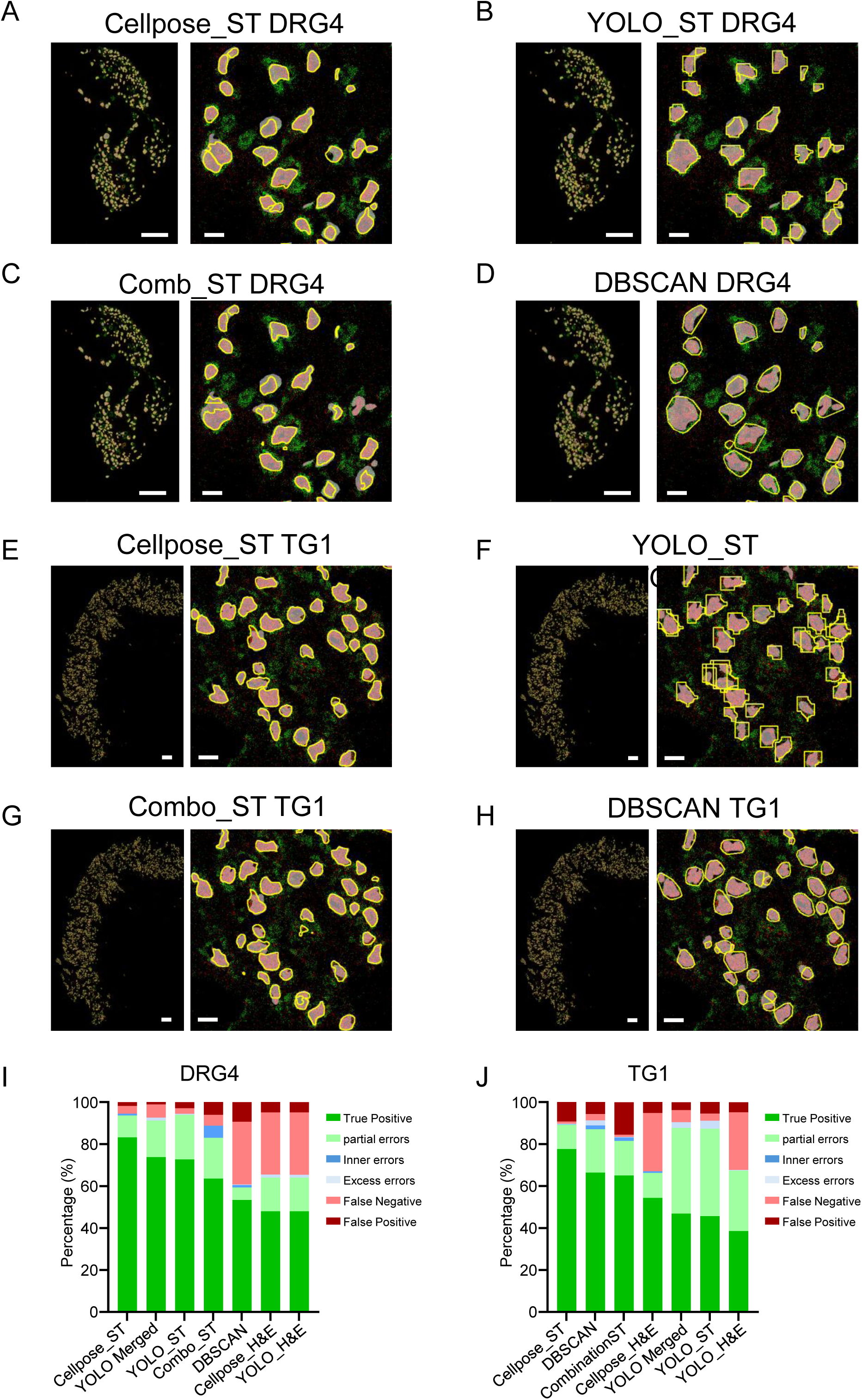
Model comparison on test datasets. (**A-D**) Representative segmentation results of each model’s predictions on the DRG4 test dataset: Cellpose_ST (A), YOLO_ST (B), Combo_ST (C) and DBSCAN (D). Predicted neuron boundaries (yellow) overlaid on ST image. Scale bars: 500 µm in (full image) and 50 µm in (zoomed-in image). (**E-H**) Representative segmentation results of each model’s predictions on the TG1 test dataset: Cellpose_ST (E), YOLO_ST (F), Combo_ST (G) and DBSCAN (H). Predicted neuron boundaries (yellow) overlaid on ST image. Scale bars: 500 µm in (full image) and 50 µm in (zoomed-in image). (**I**-**J**) Summary of all models’ performance on the test datasets DRG4 (I) and TG1 (J).

To quantitatively compare segmentation performance, we calculated different metrics (Supplementary Table 1) and categorized prediction outcomes as different categories (Figure 6I-J). For DRG4, Cellpose_ST achieved the highest F1-50 score and the largest proportion of true positive predictions. The YOLO_Merged model performed below Cellpose_ST, yet it surpassed YOLO_ST slightly. Combo_ST and DBSCAN showed moderate performance with lower true positive rates, while the H&E-only models, Cellpose_H&E and YOLO_H&E, exhibited worse performance. For TG1, Cellpose_ST consistently remained the top performer. DBSCAN and Comb_ST achieved reasonable results but were slightly less accurate. YOLO_Merged slightly outperformed YOLO_ST again.

### Downstream analysis is sensitive to segmentation accuracy

To evaluate whether sensory neuron segmentations from different models are suitable for downstream analyses and applications, we applied a pipeline to quantify transcript counts for each gene within segmented cells and generated a gene expression matrix (see Methods and Github link: https://github.com/taimeimiaole/Xenium-Segmentation). This matrix was then used for cell clustering, marker gene identification, and other transcriptomic analyses. Cell clustering was performed using Seurat^29^ based on segmentations from various models. All models produced a similar number of clusters compared to manual annotation (Figure S10A–E).

We selected DBSCAN, Cellpose_ST, Combo_ST, and YOLO_ST for direct comparison with manual annotations through co-clustering. In UMAP space, cells from these models were generally well intermingled with manually annotated cells (Figures 7A–B & S12A-B). Cluster correlation analyses further revealed high concordance overall (Figures 7C–D & S12C-D), but notable differences emerged. For example, manual Cluster 1 was split into two clusters in Cellpose_ST, DBSCAN, and YOLO_ST, yet remained a clean one-to-one match in Combo_ST. Conversely, Cluster 9 split in Combo_ST but not in the other three models. Cluster 6 was divided into two subclusters by DBSCAN and into three by YOLO_ST, while remaining intact in Cellpose_ST and Combo_ST. Cluster 14—lacking clear marker genes in the original publication^26^— was absent from all automated segmentations, and Cluster 13 was completely missed by YOLO_ST. These discrepancies underscore that segmentation accuracy directly impacts the resolution and fidelity of cell type identification.

**Figure 7.**
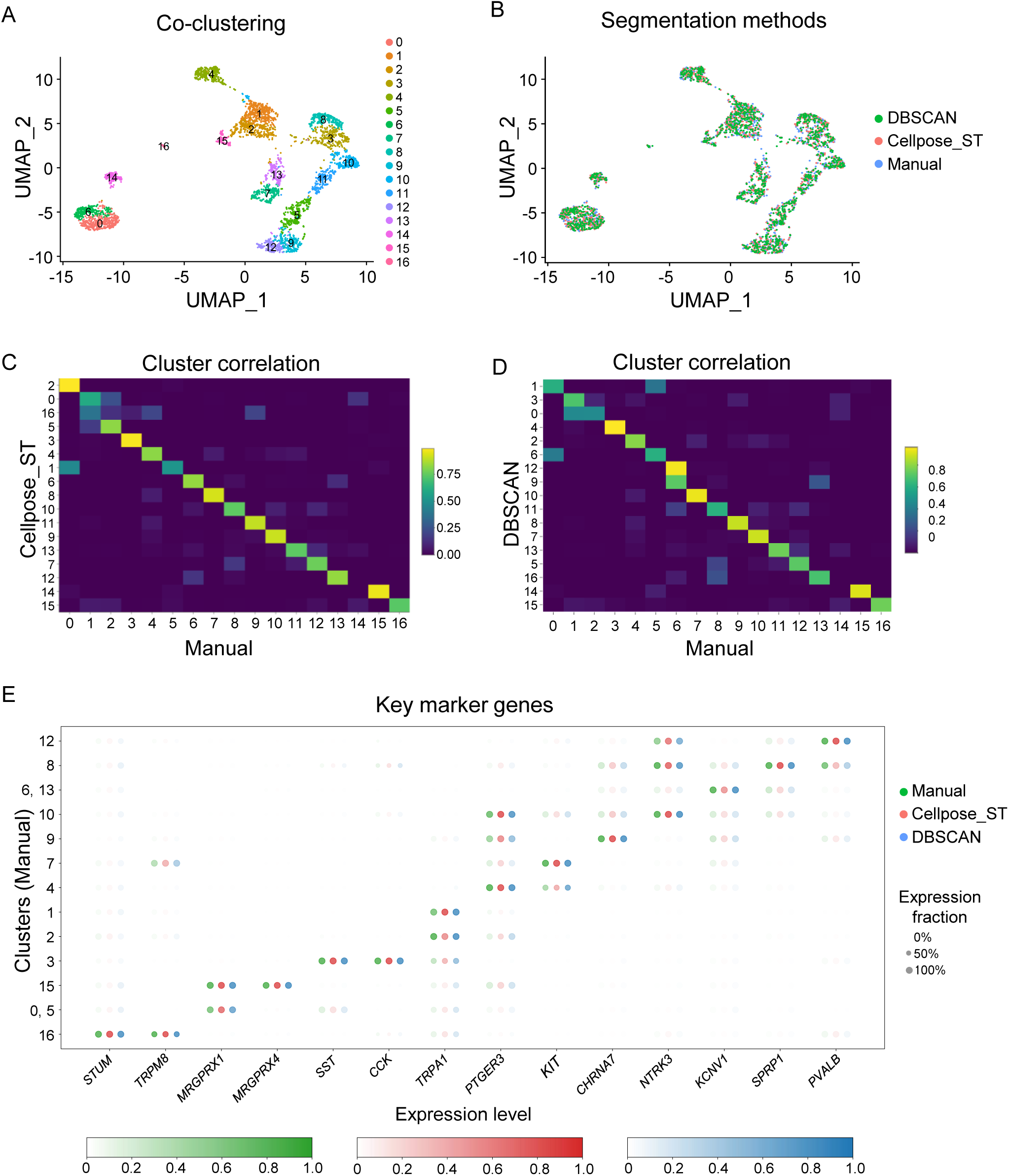
Model comparison with downstream analysis. (**A**) UMAP plot showing the co-clustering of cell segmentation from manual annotation, DBSCAN model and Cellpose_ST model. (**B**) UMAP plot showing the distribution of cell segmentation from manual annotation, DBSCAN model and Cellpose_ST model in each cluster. (**C-D**) Heatmap comparing the similarities of cell clusters between Cellpose_ST (C) or DBSCAN (D) segmentation and manual annotation. (**E**) Dot plot showing the top marker genes expressed in each cell cluster from manual annotation (green), Cellpose_ST (red), and DBSCAN (blue). The color scale represents the normalized average expression level (from 0 to 1) of each gene in the clusters.

For most clusters, key marker genes identified from DBSCAN, Cellpose_ST, YOLO_ST, and Combo_ST closely matched those from the corresponding manual clusters (Figures 7E, S11, S12E). One exception was Cluster 10 in YOLO_ST, where the expected marker NTRK3 was missed entirely.

### A quality check step using manual review and correction

The discrepancies observed in downstream analyses revealed a key point: even well-performing models can produce segmentation errors that alter outcomes and conclusions. To minimize this concern, we implemented a quality check step by adding manual review and correction of automated segmentation results.

This process was implemented using the ROI Manager in ImageJ/FIJI. Because ROI Manager uses the .roi file format to store segmentation masks (or regions of interest, ROIs), we first created a conversion pipeline to support a variety of model output formats, including .txt, .csv, and .npy, by converting them into the .roi format compatible with ROI Manager (see Methods and Github link: https://github.com/taimeimiaole/Xenium-Segmentation). Corrections followed four straightforward steps: loading segmentations, identifying errors, editing incorrect segmentations, and saving the updated masks (Figure 8A). Corrections were classified into three categories: “add” for missed neurons (false negatives), “delete” for incorrect predictions (false positives), and “edit” for inaccurate boundaries—typically requiring deletion of the original mask and redrawing the boundary using the ROI Manager’s freehand tool.

**Figure 8.**
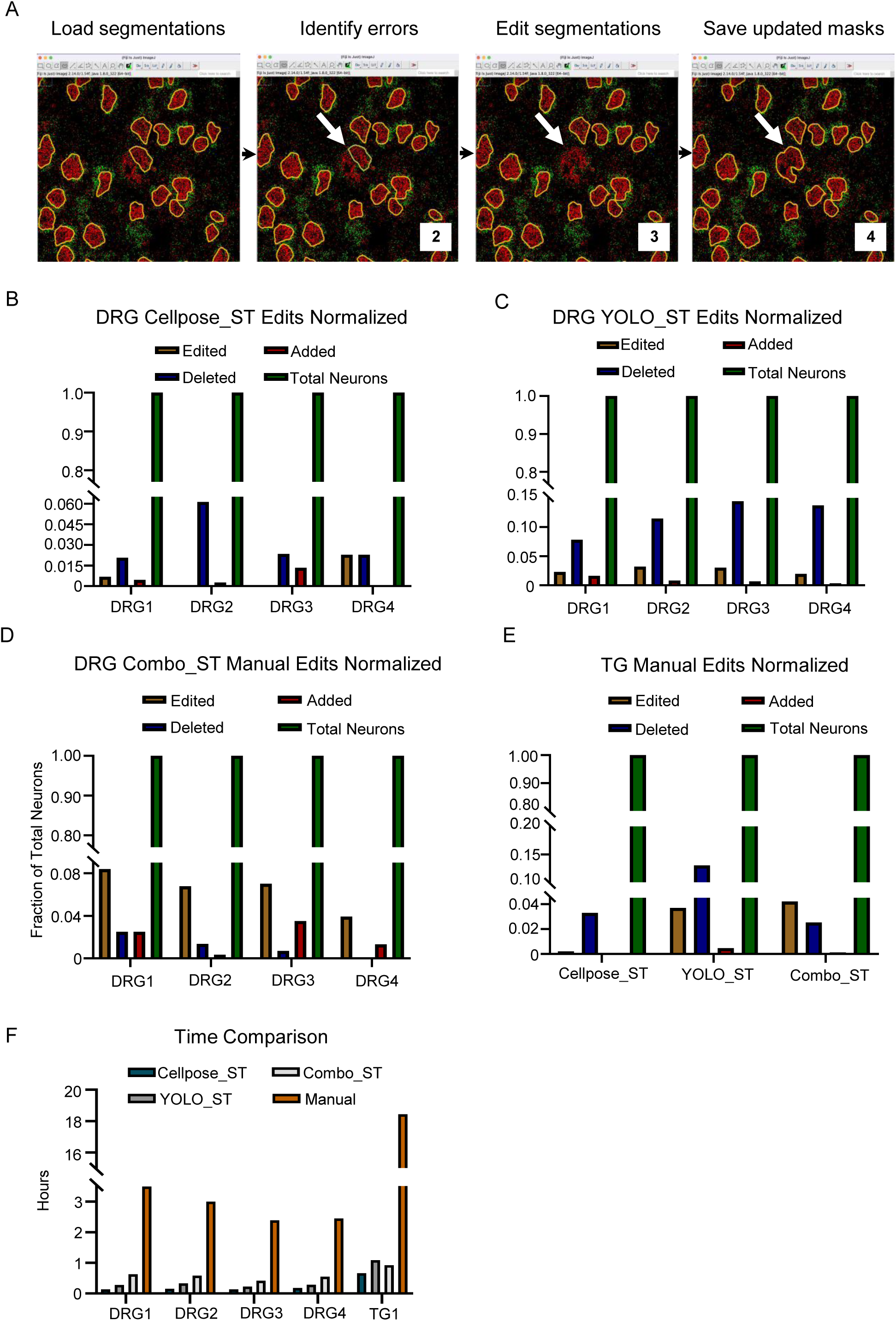
Manual review and correction pipeline for segmentation results. (**A**) Schematic of the manual correction process using the ImageJ/FIJI ROI Manager and the Freehand Selection tool. (**B-D**) Quantification of manual edits on DRG1–DRG4 segmentations for three models: Cellpose_ST model (B), YOLO_ST (C) and Combo_ST (D). Edits are categorized as Edited (modifying boundaries), Added (missed neurons), or Deleted (false positives), and are normalized to the total number of segmented neurons. (**E**) Manual correction summary for TG1 segmentation using the same edit categories. (**F**) Time comparison between manual correction following each segmentation method and manual-only segmentation.

Manual correction was applied to predictions from the Cellpose_ST, YOLO_ST, and Combo_ST models across DRG1–4 and TG1. Among them, Cellpose_ST required the fewest corrections, primarily involving deletion of false-positive neurons (Figure 8B). In contrast, YOLO_ST required more edits (Figure 8C), including both false-positive deletions and boundary refinements, due to the boxy shape of some predictions. Combo_ST fell in between, requiring fewer corrections than YOLO_ST but more than Cellpose_ST (Figure 8D). For TG1, all three models required corrections for only a small fraction of the predicted neurons (Figure 8E), with Cellpose_ST again needing the least correction, followed by Combo_ST and YOLO_ST.

Combining automatic prediction with manual correction significantly reduced the total segmentation time and effort when compared to manual segmentation (Figure 8F). Compared to the 2–3 hours typically needed for full manual segmentation of a DRG section, correcting Cellpose_ST predictions took less than 10 minutes per section on average. For TG1, the correction time was approximately 40 minutes, while the original manual segmentation time was more than 18 hours. Significant time was also saved by YOLO_ST and Combo_ST, though correction time was longer than time needed for Cellpose due to their higher error rates. These results highlight the efficiency and practicality of this hybrid approach—automated segmentation followed by manual review and correction improves both speed and accuracy in cell segmentation workflows of spatial transcriptomics analysis.

## Discussion

Accurate cell segmentation remains one of the most consequential—but often underappreciated—determinants of reliability in analyzing probe-based spatial transcriptomics datasets. Using human sensory ganglia (DRG and TG) as a representative example, we systematically evaluated multiple segmentation approaches for targeted neuronal analysis and developed a comprehensive workflow encompassing raw data preprocessing, segmentation, parameter optimization, downstream transcriptomic analysis, and manual quality control. From our experience, when performing cell segmentation of probe-based spatial transcriptomics, it is important to consider that : (1) marker genes should include those facilitating cell boundary delineation; (2) method choice depends on targeted cell types and tissues; (3) parameter tuning and optimization are necessary for every method; (4) all algorithms exhibit some characteristic errors that will influence downstream analysis and outcomes; and (5) manual review will improve accuracy of automatic models.

### Probe panel design for optimal segmentation

Regardless of whether a clustering-based or deep learning approach is chosen, segmentation fundamentally depends on the quality of marker gene transcript data. For cell types of interest, selecting appropriate markers is therefore critical. It is desirable to include 2–4 candidate cell type–specific markers in the probe panel, as marker quality (e.g., signal-to-noise ratio) may vary substantially. Multiple candidates increase the likelihood of obtaining at least one high-quality signal, and this foresight during experimental design will help segmentation performance. Another consideration is that some markers should clearly delineate boundaries of targeted cells, such as UCHL1 and FABP7 used in this study.

### Segmentation method choice

Method selection should be guided by the biological question, annotation resources, and tissue architecture. Deep learning approaches can achieve high accuracy when trained on sufficiently large, well-annotated datasets, but the need for manual labeling is a common bottleneck. Clustering-based methods such as DBSCAN require minimal labeled data and, when parameters are well tuned, can perform comparably to supervised models, making them especially useful when annotation is not available. In tissues with densely packed cells and indistinct boundaries (e.g., spinal cord interneurons), accurate manual annotation is impractical. In such cases, clustering-based methods may be a better choice than deep learning approaches. Computational constraints, such as GPU memory, are another important consideration in method selection.

### Parameter tuning and optimization

Across both clustering-based and deep learning methods, parameter optimization is a primary driver of segmentation accuracy. In clustering-based approaches such as DBSCAN, optimal parameters are closely linked to target cell size and transcript density. In deep learning models, parameters such as the number of training epochs and the size of the training set require systematic evaluation. Each architecture introduces additional, method-specific sensitivities: for example, in Cellpose, the minimum mask threshold—which controls whether an image is used in training—can substantially affect performance. In YOLO, the max_det parameter caps the number of predicted objects per image; setting it too low misses detections. These examples underscore that segmentation performance emerges from the interplay of algorithm design, parameter settings, and dataset characteristics—without careful tuning, even state-of-the-art methods can produce misleading results.

### Error types and their implications

No current segmentation method is perfect. Each displays characteristic error patterns. In our comparisons, Cellpose models produced smooth boundaries with relatively few errors; Combo models generated accurate boundaries but more false negatives; YOLO models achieved high detection rates but yielded box-like boundaries; DBSCAN was highly parameter-sensitive and often produced slightly oversized boundaries. Segmentation errors can be categorized as false positives, false negatives, and boundary imperfections (including partial matches, inner, or excess, as discussed previously). We speculate that these error modes have distinct impacts on downstream analyses: boundary imperfections may incorporate transcripts from neighboring cells, blurring cluster boundaries and leading to cell type misclassification; false positives may introduce spurious cell types; and extensive false negatives may result in the omission of rare populations. These hypotheses could be systematically tested in future studies to quantify how segmentation errors propagate into clustering outcomes and cell type assignments.

### Manual review and correction

While automated models can generate rapid and reasonably accurate segmentations, our results suggest that manual review and correction is a good step to have. To facilitate this, we developed an easy-to-adopt pipeline for manual inspection and refinement, compatible with diverse model output formats. This step is not merely cosmetic—undetected segmentation errors can propagate into downstream analyses, leading to misclassification of cell types, loss of rare populations, or erroneous marker gene assignments. In our view, incorporating a quality-control stage is an essential step for drawing reliable biological conclusions from spatial transcriptomics data.

### Future directions

Looking ahead, we see several promising avenues for improving segmentation accuracy. First, more effective integration of H&E or other histological images—implemented through multimodal architectures capable of processing multi-channel inputs during training and inference—may enhance recognition of complex cell morphologies. Second, incorporating additional imaging modalities, such as dyes or markers labeling cell membranes, cytoplasm, and nuclei, could provide complementary structural cues that improve model robustness. Third, testing additional deep learning architectures, as they are rapidly evolving and advancing, may further enhance accuracy, stability, and generalizability.

### Consortia

The members of the Penn High Precision Pain Center are Zarina S. Ali, Maria Bograkou, Caitlin Cronin, Patrik Ernfors, Charles Hwang, Ewa Jarocka, Eric A. Kaiser, Faiz Kassim, Åsa Rydmark Kersley, Katarina Laurell, Julie Leu, Mingyao Li, Ying Li, Dongming Liang, Wenqin Luo, Anne Marshall, Lotta Medling, Saad Nagi, Johan Nikesjö, Håkan Olausson, Dorota Persson, Juan Inclan Rico, Ebenezer Simpson, Oumie Thorell, Frida Larsson Torri, Dmitry Usoskin, Hao Wu, Hanying Yan, Huasheng Yu, Eric Zager.

## Acknowledgements

We would like to thank the donors and their families for the precious gifts that are fundamental for this study. We thank the National Disease Research Interchange (NDRI) for procuring human DRG samples; the National Institute of Mental Health (NIMH) Human Brain Collection Core (HBCC) for providing human TG samples; Drs. Stefano Marenco and Yiyin Liu in NIMH-HBCC for their help and support; the Molecular Pathology and Imaging Core (MPIC) for providing the platform and support for conducting Xenium spatial transcriptomics. We also thank Sansan Lu and Ava Girshovich, two Harriton High School, PA, students, for their assistance with manual segmentation of hTG tissues, as well as Shiwei Chen for setting up the remote-control system. Drs. Wenqin Luo, Mingyao Li, Penn HPPC, and this research are supported by a National Institutes of Health (NIH) HEAL U19 grant (U19-NS-135528). Dr. Wenqin Luo is also supported by NIH grants R01-NS-131209, U01-EY-034681 and R21-NS-142656. Mingyao Li is also supported by R01HG013185 and R01LM014592.

## Author Contributions Statement

H.Yu, M.L., and W.L., conceptualized the project. N.W. worked on dataset curation, development of the Baysor model, and creation of the conversion pipeline. H.Yan developed the DBSCAN model and optimization. A.C. developed the YOLO model, the Combination Model, merged images, and manual correction pipeline, as well as completing manual segmentation of TG1 and performing manual correction. K.S. developed the Cellpose model, model evaluation pipeline, and stitching pipeline. H.Yu generated the Xenium data, performed downstream analyses, and worked closely with N.W., H.Yan, A.C., and K.S. to develop and optimize each model. H.Yu, A.C., K.S., N.W. and H.Yan wrote the manuscript draft. W.L. and M.Y. reviewed the manuscript.

## Declaration of Interests

W.L. has received research grants from Eli Lilly, and her spouse is an Eli Lilly employee and holds equity. H.Yu, M.G., and W.L. are co-founders of Nociheal LLC and hold equity of the company. M.L. is a co-founder of OmicPath AI and hold equity of the company.

Figure Generation Software

Figures were generated in Powerpoint (Microsoft Office) and GraphPad Prism (v8 & v9, GraphPad Software Inc. La Jolla, CA, USA). Some elements in the figures were created with assistance of ChatGPT, including the Baysor, DBSCAN, Cellpose, and YOLO symbols (Figures 2A, 3A, 4 A, S4A, S6A) as well as part of the cartoon in Figure 1A.

## Data Availability

The raw and processed datasets for the 10x Xenium transcriptomics of human DRG and TG neurons are available on Dryad. The dataset link is currently private for peer review and will be made public upon acceptance of the manuscript for publication.

## Code Availability

The custom code used in this study is available on GitHub at https://github.com/taimeimiaole/Xenium-Segmentation.

## Methods

### Spatial transcriptomics and H&E staining

Spatial transcriptomics was performed using the Xenium Analyzer (10x Genomics) following the manufacturer’s instructions (https://cdn.10xgenomics.com/image/upload/v1710785024/CG000584_Xenium_Analyzer_User Guide_RevE.pdf). In brief, 10-μm-thick tissue cryosections were mounted on Xenium slides, mRNAs were targeted using a custom gene panel consisting of 100 genes, including 87 neuronal marker genes and 13 non-neuronal marker genes^26^. Post-Xenium H&E staining was subsequently performed. H&E staining images were acquired using the Aperio Scanner microscope system.

### Baysor segmentation and evaluation

Baysor runs were carried out following the guidelines described in the original publication^13^. For every run, a dedicated folder containing a custom.yaml file was created to define the parameter constraints. Segmentation was then executed through a sequence of Julia prompts. Segmentation results were saved in geojson format for downstream analysis. The custom.yaml file included parameters such as *exclude_genes*, *scale*, *scale_std*, *min_molecules_per_cell*, *n_clusters*, and *iters*. In this study, we primarily tested two parameters: *min_molecules_per_cell* (reflecting transcript signal density and cell size) and *scale* (related to cell size). The *scale_std* was adjusted to 15%. The *prior segmentation confidence* was set to 0.0, and the *number of iterations* was fixed at 500, although convergence was typically reached before this limit. Other parameters were generally left at their default settings; for example, *min_molecules_per_segment* and *confidence_nn_id* were automatically determined by the value of *min_molecules_per_cell*. For the evaluation, segmentation outputs in geojson format were first converted to CSV files to extract cell boundary outlines, and subsequently to ROI files for visualization in ImageJ. This enabled direct comparison between predicted and manual masks on both the original spatial transcriptomics (ST) image and the corresponding H&E stain. The evaluation metrics were calculated by evaluation pipeline.

### DBSCAN segmentation and evaluation

For DBSCAN prediction, a cluster-based cell segmentation pipeline was developed to import Xenium transcript outputs into R^30^. DBSCAN clustering^31^ was applied to spatial coordinates of UCHL1, the selected pan-neuronal marker, to group neuronal transcripts. For each cluster, a convex hull^32^ was computed to define the neuron boundary. The optimal range of DBSCAN parameters Eps and MinPts were estimated using our proposed formula. In the formula, the cell size (r) and the transcript density (d) of each neuron were calculated as 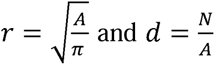, where N is the number of neuronal transcripts and A is the area of a specific neuron. For each chosen Eps value, it is not necessary to test every MinPts integer value. Larger Eps values allow for larger MinPts, so wider step sizes can be used.

### ST image generation and H&E image alignment

ST images were generated from the Xenium transcript CSV file, with different transcripts displayed in pseudocolor. The image scale was set according to the ratio of the Xenium capture area, which was determined from the nucleus staining images. The resolution of ST images was set to 1 pixel per micrometer, and each transcript coordinate corresponded to 1 pixel. H&E images were then aligned to the ST images by matching the corresponding nuclear staining patterns between the Xenium and H&E images. To ensure comparability, H&E images were downsampled to the same resolution as the ST images. Finally, the images were overlaid from the top-left corner to verify proper alignment.

### Manual cell segmentation of DRG or TG neurons

Manual cell segmentation was performed on ST images using ImageJ software. The ST images used for manual annotation included the pan-neuronal marker UCHL1 and the satellite glial cell marker FABP7, which helped delineate neuronal boundaries. These images were loaded into ImageJ and annotated manually with the freehand selection tool and ROI Manager. In addition, the H&E image was loaded in Xenium, overlaid with UCHL1 signals, and used as a reference to double-check the annotations.

### Training dataset generation

The training dataset for deep learning models was generated from ST images (DRG1–3) and their corresponding aligned H&E images. ST images included the pan-neuronal marker UCHL1 (pseudocolored red) and the satellite glial cell marker FABP7 (pseudocolored green). Labels for neurons were derived from manual annotations for each tissue image. Both ST and H&E images were cropped into 500 × 500 µm/pixel patches, and a corresponding label text file was generated to indicate the boundaries of each cell. For cells located at the image borders, the labels were readjusted by the pipeline’s auto-adjust function to fit the new cell shape after cropping. The cropped images were split into 80/10/10 proportions for training, validation, and testing.

### Merged image generation

To generate the merged dataset combining both ST and H&E images for training, the H&E images were first inverted and split into red, green, and blue single-channel images. Inversion was necessary to prevent the white background of the original H&E images from dominating during channel splitting. Among the channels, the red channel provided the clearest distinction of neuronal structures; this channel was converted to grayscale for subsequent RGB merging. The grayscale H&E red channel image, together with the UCHL1 ST image and FABP7 ST image (both grayscale), were then merged into RGB images, with the H&E stain assigned to the red channel, UCHL1 to the blue channel, and FABP7 to the green channel. The resulting merged images were split into training, validation, and testing sets using the same procedure applied to the ST and H&E datasets, ensuring that corresponding images were placed in the same subset across all three datasets (ST, H&E, and merged).

### Pipeline for conversion between different segmentation formats

Segmentation outputs from different models were generated in varying formats: Baysor (.geojson), DBSCAN and YOLO (.txt), Cellpose and the Combo model (.npy), and manual annotations (.roi). To enable unified evaluation and manual correction in FIJI, we developed a conversion pipeline with CSV as the intermediate format. This pipeline supported conversions from .csv to .roi, .txt, .geojson, and .npy, as well as from each of these formats back to .csv. For model evaluation, all model outputs and manual annotations were converted to .csv for use in the evaluation pipeline. For manual correction, model predictions were converted from .csv into .roi files and then packaged as a single zipped .roi file compatible with FIJI’s ROI Manager. This conversion framework enabled consistent comparison across models and seamless integration with our evaluation and correction pipelines.

### Evaluation metrics

Each segmentation method included its own in-training metrics; however, because these varied across tools, we developed a standardized evaluation pipeline (Figure S3A). Segmentation outputs were first converted into .csv format using our conversion pipeline to allow uniform downstream evaluation. Model performance was primarily assessed using the Intersection over Union (IoU) between predicted and ground truth masks, defined as the ratio of their intersection to their union. Since IoU alone does not account for false negatives or false positives, we additionally calculated precision, recall, and F1-score at an IoU threshold of 0.5, or at thresholds from 0.5 to 0.95. Pixel-weighted precision, recall, and F1-scores were also included in the evaluation metrics. In total, the pipeline reported nine main statistics: cell-weighted precision, recall, and F1 at IoU = 0.5; average precision, recall, and F1 across IoU thresholds 0.5–0.95; and pixel-weighted precision, recall, and F1. Beyond numerical metrics, the pipeline also categorized segmentation errors based on their overlap with ground truth masks (Figure S3B). Errors were classified as true positives (IoU > 0.5), partial errors (partial overlap, IoU ≤ 0.5), inner errors (predicted mask entirely inside the ground truth, IoU ≤ 0.5), excess errors (predicted mask larger than the ground truth, IoU ≤ 0.5), false negatives (missing prediction), and false positives (prediction without a corresponding ground truth).

### YOLO training and evaluation

The YOLO (Version 8) model was trained on three-channel images with corresponding annotations organized into train, test, and validation subfolders, as specified in a dataset .yaml file. We used the small-sized YOLO segmentation model as the backbone for further training. All YOLO models are pretrained on either Microsoft’s COCO dataset or Imagenet. Training parameters were optimized for our dataset by adjusting the model size and number of epochs, and by disabling mask overlap (overlap_masks=False). Training automatically included validation after each epoch and a final validation upon training completion. It is critical to specify the image size during both training and prediction, as YOLO otherwise resizes images to 640×640 pixels by default, which can impair accuracy. During training, additional outputs such as learning curves can be saved (save_plots=True). The trained model is saved in PyTorch format (.pt), with two checkpoints: best.pt (model saved at highest-performing iteration) and last.pt (model saved at final training iteration). The two models may give different outputs. For prediction, the trained model path, image size, and additional parameters must be specified. In our workflow, bounding boxes were suppressed (to display only segmentation masks), and annotations were saved as text files (save_txt=True). Annotation files are automatically named to match their corresponding images. The final YOLO models—YOLO_ST, YOLO_H&E, and YOLO_merged—were trained with the following key parameters: epochs = 100, overlap_masks = False, image size = 640.

### Cellpose training and evaluation

To enable direct comparison across deep-learning models, the same training, validation, and test datasets were used. However, because Cellpose does not support validation during training, the validation dataset was excluded, and the model was trained and evaluated using the training and test datasets only. Model optimization focused on key parameters, including pre-trained model selection (model), channel specification (c1, c2), number of epochs (n_epochs), minimum masks per image (min_train_masks), and flow and probability thresholds (flow_threshold, cellprob_threshold). A full list of Cellpose training parameters can be found in the Cellpose API guide (https://cellpose.readthedocs.io/en/latest/api.html). At each optimization step, parameters under evaluation were varied while all others were held at default settings, and performance was assessed with the standardized evaluation pipeline. Ground-truth annotations were formatted for Cellpose by generating .npy mask files from the manually annotated mask directories, with filenames corresponding to each image plus the suffix _masks.npy. To ensure compatibility with the min_train_masks threshold, images without annotated neurons were removed prior to training. Training inputs included the training image directory, the test image directory, and the specified parameter set. Cellpose outputs include segmentation arrays (_seg.npy), flow maps, and colored masks. We additionally enabled the .roi file export to generate .csv files for downstream evaluation. For prediction, the cellpose.eval method was applied to individual images using the optimized model. The model.eval call returned masks, flows, and styles, and segmentation masks were also saved as ROIs using cellpose.io.save_rois for integration into the manual correction pipeline. Because Cellpose performance degrades in the presence of large empty regions, input images were cropped to the tissue area prior to prediction.

### Combination model training and evaluation

The Segmentation Model’s Pytorch modules (Version 0.3.4) included nine architectures and ten encoders. Our goal was to identify the best architecture–encoder combination. For each run, the dataset path and number of epochs were specified, and a validation mAP50 score was generated after each epoch. Models were trained for 80 epochs. Not all architectures were compatible with all encoders, and some combinations failed due to excessive GPU or memory requirements. These combinations were discarded. To systematically evaluate performance, we first trained each architecture with the smallest encoder from each encoder family (“first combinations”), yielding 72 valid combinations. Average mAP50 scores across the last five epochs were used to assess accuracy. From this analysis, we identified the top three architectures and encoders, although UNet++—despite performing well—was excluded due to incompatibility with the top-performing encoders. Next, we trained the top three architectures with each of the sizes of the top three encoders (“second combinations”), resulting in 42 combinations. As before, the overall score for each combination was defined as the average mAP50 across the last five epochs. From these, we selected the three best-performing combinations. These top combinations were then trained with varying epoch numbers (5, 10, 20, 35, 50, 75, 100, 200, 300) to determine the optimal training epoch. The UNet + VGG16_bn combination was most consistent in its accuracy across epochs and was selected as the final model. This model was subsequently trained and tested for comparison with the other models described in this study. For prediction, tissue images were cropped into 500 × 500 pixel tiles and placed in the test/images subfolder. A corresponding test/labels subfolder was required, though it remained empty. Predictions were generated using a threshold of 0.8 and saved as .npy files containing binary and probability arrays for each tile. A custom pipeline was then applied to merge the tile arrays into a single array representing the full tissue, which was converted into .csv format for downstream evaluation.

### Manual correction

Manual correction of model outputs was performed in ImageJ/FIJI. Predictions from YOLO, Cellpose, and Combo model were first converted into zipped .roi files using our custom format-conversion pipeline. Each model’s predictions were corrected independently. For each tissue, the ST image (UCHL1+FABP7) was loaded into FIJI, and the corresponding .roi file was imported into the ROI Manager. Predicted annotations were reviewed and corrected according to three error categories: (1) addition of annotations to address false negatives, (2) deletion of annotations to remove false positives, and (3) editing of annotations to correct boundary errors. Incorrect predictions were deleted by selecting and deleting the ROIs, while additions and edits were made using FIJI’s freehand selection tool and added to the ROI Manager. Masks were edited only if they contained approximately less than half of the expected transcripts by eye; masks estimated to capture at least three-quarters of transcripts were retained without modification. H&E images of the same tissue were used as a reference to guide corrections, providing additional information about neuronal presence and boundaries that were sometimes ambiguous in the ST images. Once corrections were complete, the finalized annotations were saved as zipped .roi files from FIJI.

---

**Figure S1.**
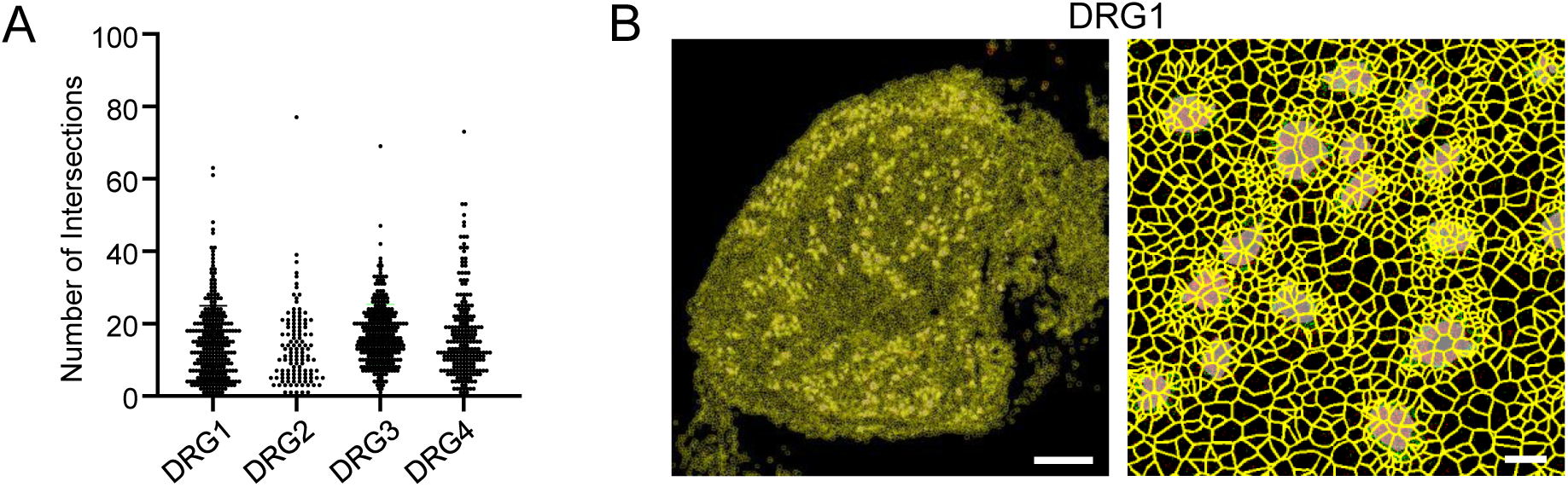
Xenium automated cell segmentation. (**A**) Quantification showing the number of fragments, which was split into by the Xenium automated cell segmentation, for each manually annotated neuron in DRG1–4. Each dot represents a single manually segmented neuron. (**B**) Example segmentation results from the Xenium automated segmentation for DRG1: a view of the entire tissue section (left), a zoomed-in view with segmentation boundaries (yellow outline) overlaid with ST images (right). Scale bars: 500 µm in (left) and 50 µm in (right).

**Figure S2.**
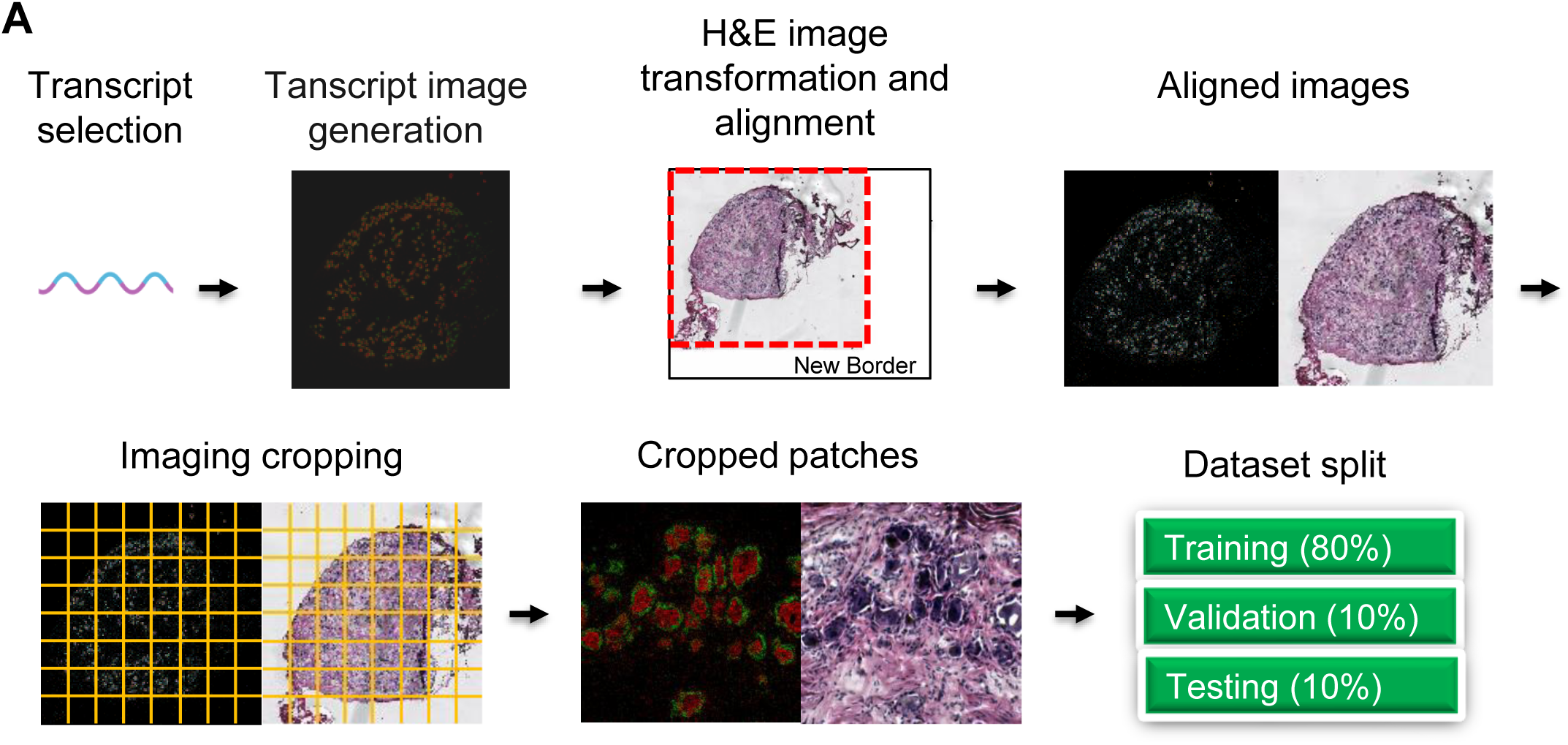
Dataset preparation for manual segmentation and deep learning model training. (**A**) Workflow for generating the training dataset used for YOLO, Cellpose, and other deep learning segmentation models. Steps include transcript selection, ST image generation, H&E image transformation and alignment, cropping the ST and corresponding H&E images to generate uniform patches, and dataset split of the cropped patches into training (80%), validation (10%), and testing (10%) sets.

**Figure S3.**
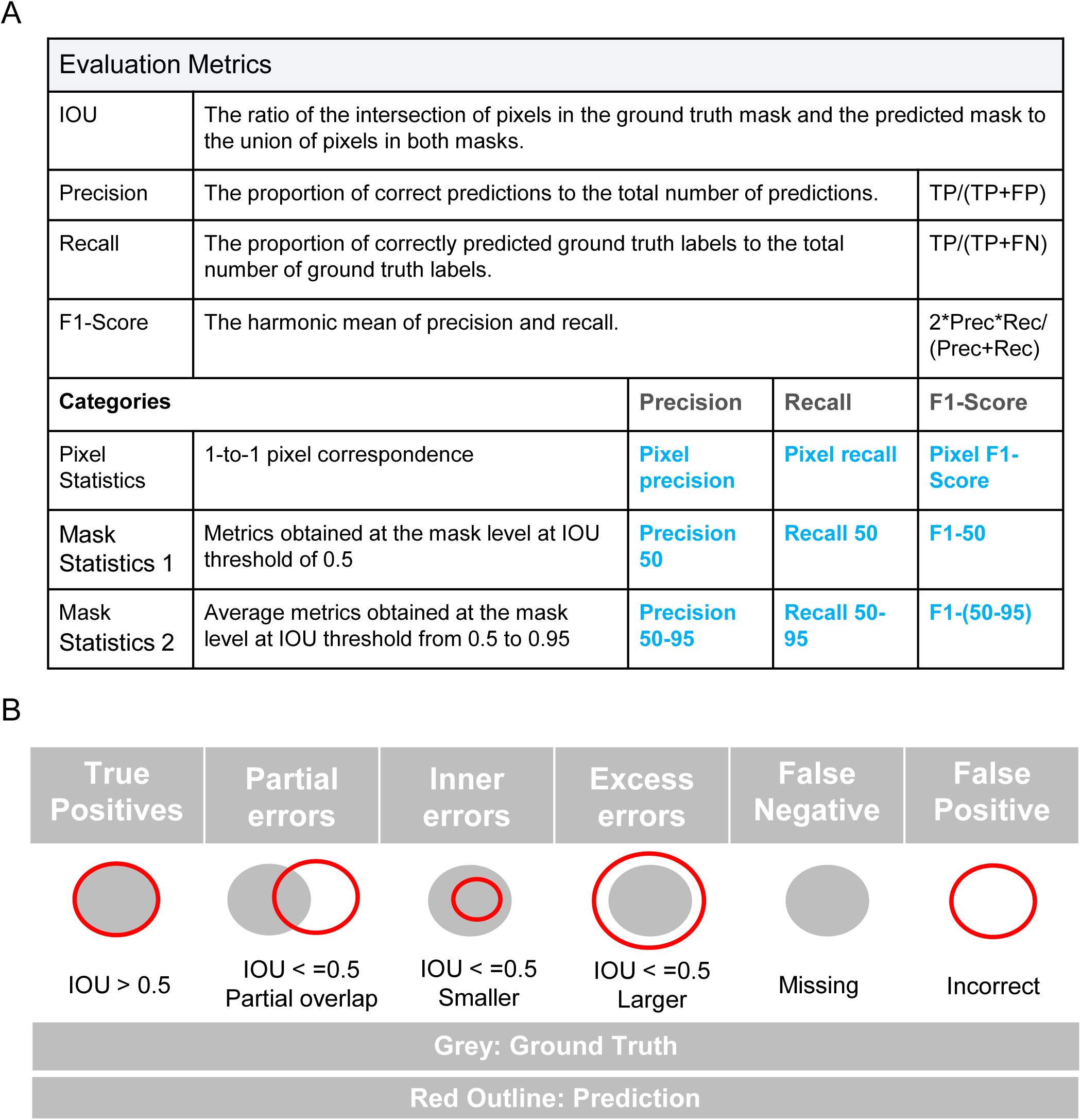
Evaluation metrics. (**A**) Summary of evaluation metrics used to assess segmentation performance. Definitions for IOU (intersection over union), precision, recall, and F1-score are provided along with the specific thresholds and categories used for analysis, including pixel-level and mask-level statistics. (**B**) Illustration of segmentation categories. Predictions are compared against ground truth masks and categorized as True Positives (IOU > 0.5), Partial errors (IOU ≤ 0.5, partial overlap), Inner errors (IOU ≤ 0.5, prediction smaller than ground truth), Excess errors (IOU ≤ 0.5, prediction larger than ground truth), False Negatives (missed detections), and False Positives (false predictions). Ground truth is shown in grey, and predictions are outlined in red.

**Figure S4.**
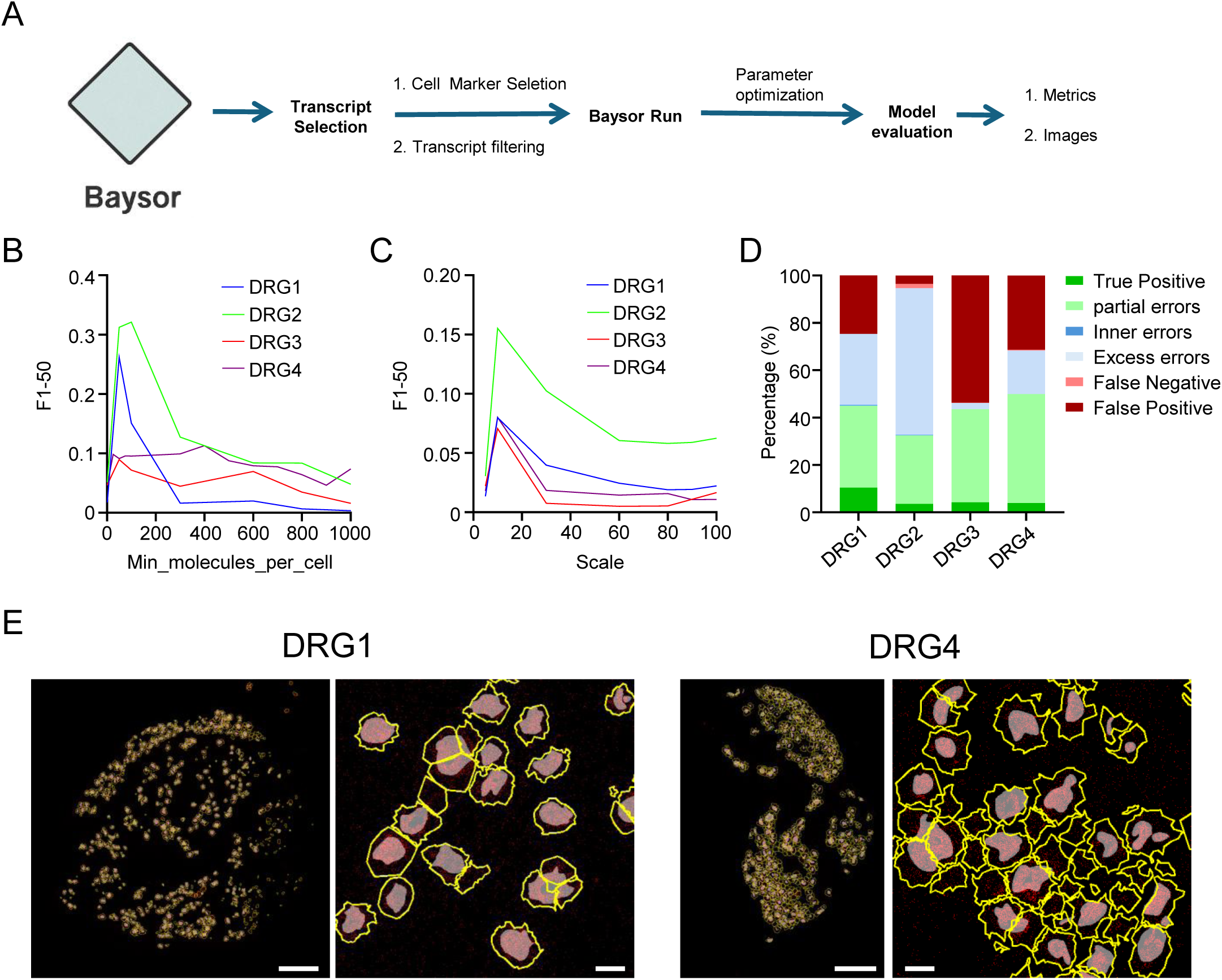
Unsupervised segmentation by Baysor. (**A**) Overview of the Baysor segmentation workflow. Key steps include transcript selection, Baysor execution with parameter optimization, and model evaluation. (**B**) F1-50 scores across varying Min Molecules (min_molecules_per_cell) with a fixed scale of 15, using UCHL1 as the neuronal marker. Results are shown for DRG1–4. (**C**) F1-50 scores across varying scale values with min_molecules_per_cell fixed at 100 and UCHL1 as the neuronal marker. Results are shown for DRG1–4. (**D**) Segmentation performance and classification of errors of Baysor’s predictions for DRG1-4. Baysor parameters were optimized specifically for each DRG when running the predictions. (**E**) Representative segmentation results for DRG1 and DRG4. Left: full tissue view; Right: 500 × 500 µm zoomed-in region showing overlay of ground truth (gray fill) and predicted boundaries (yellow outlines). Scale bars: 500 µm in full images and 50 µm in zoomed-in images.

**Figure S5.**
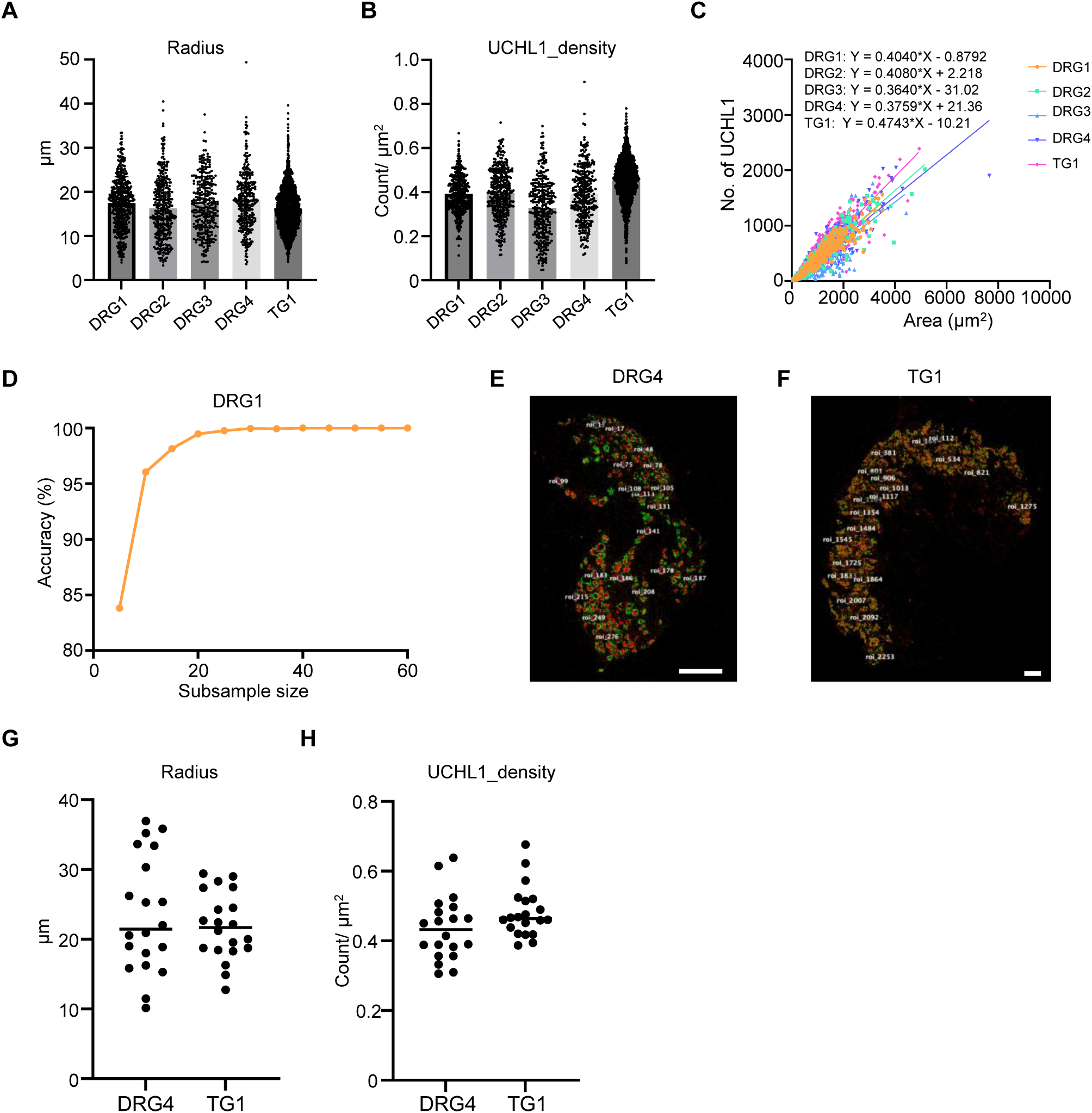
Unsupervised clustering-based segmentation by DBSCAN. (**A**) Distribution of neuronal cell size from manual annotations in DRG1–4 and TG1. (**B**) Distribution of UCHL1 transcript density (molecule count per µm²) in manually segmented neurons in DRG1–4 and TG1. (**C**) Linear regression between UCHL1 molecule count and cell size in DRG1–4 and TG1. (**D**) Sample size required to predict DBSCAN parameter Eps and MinPts. As the number of randomly selected neurons increased from 5 to 60 (x-axis), optimal Eps and MinPts range prediction accuracy (y-axis) increased. (**E–F**) Twenty neurons were randomly selected from DRG4 and TG1 to estimate optimal ranges of DBSCAN parameter Eps and MinPts. Scale bars: 500 µm. (**G–H**) Distribution of cell size and UCHL1 transcript density in 20 randomly selected neurons from DRG4 and TG1 in (E) and (F).

**Figure S6.**
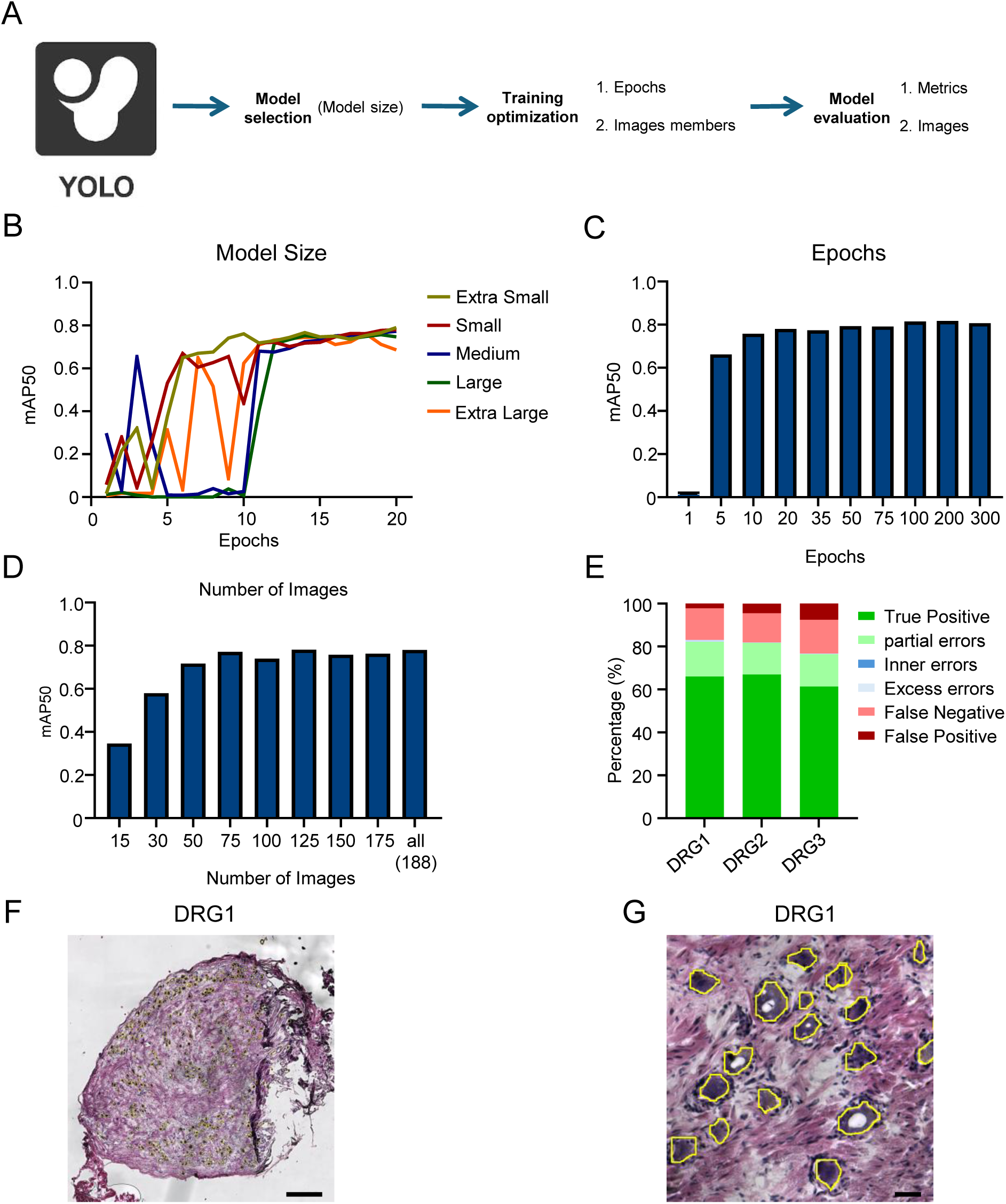
H&E image-based segmentation by YOLO. (**A**) Overview of the YOLO segmentation workflow using H&E images, including model size selection, training optimization (model size, epochs, and image set size), and evaluation. (**B**) Effect of model size on segmentation performance. mAP50 values are shown for five YOLO model sizes (from extra small to extra large) trained on DRG1–3 H&E images over 20 epochs. (**C**) Effect of training epochs on model performance. The YOLO small model was trained with 188 images for various numbers of epochs (1 to 300). (**D**) Effect of training set size. The YOLO small model was trained for 20 epochs using increasing numbers of training images. (**E**) Segmentation error classification for the YOLO_H&E model applied to DRG1–3 full tissue images. (**F–G**) Representative segmentation results for DRG1 whole image (F) and a zoomed-in area (G). Predicted neuron boundaries (yellow) overlaid on H&E image. Scale bars: 500 µm in (F) and 50 µm in (G).

**Figure S7.**
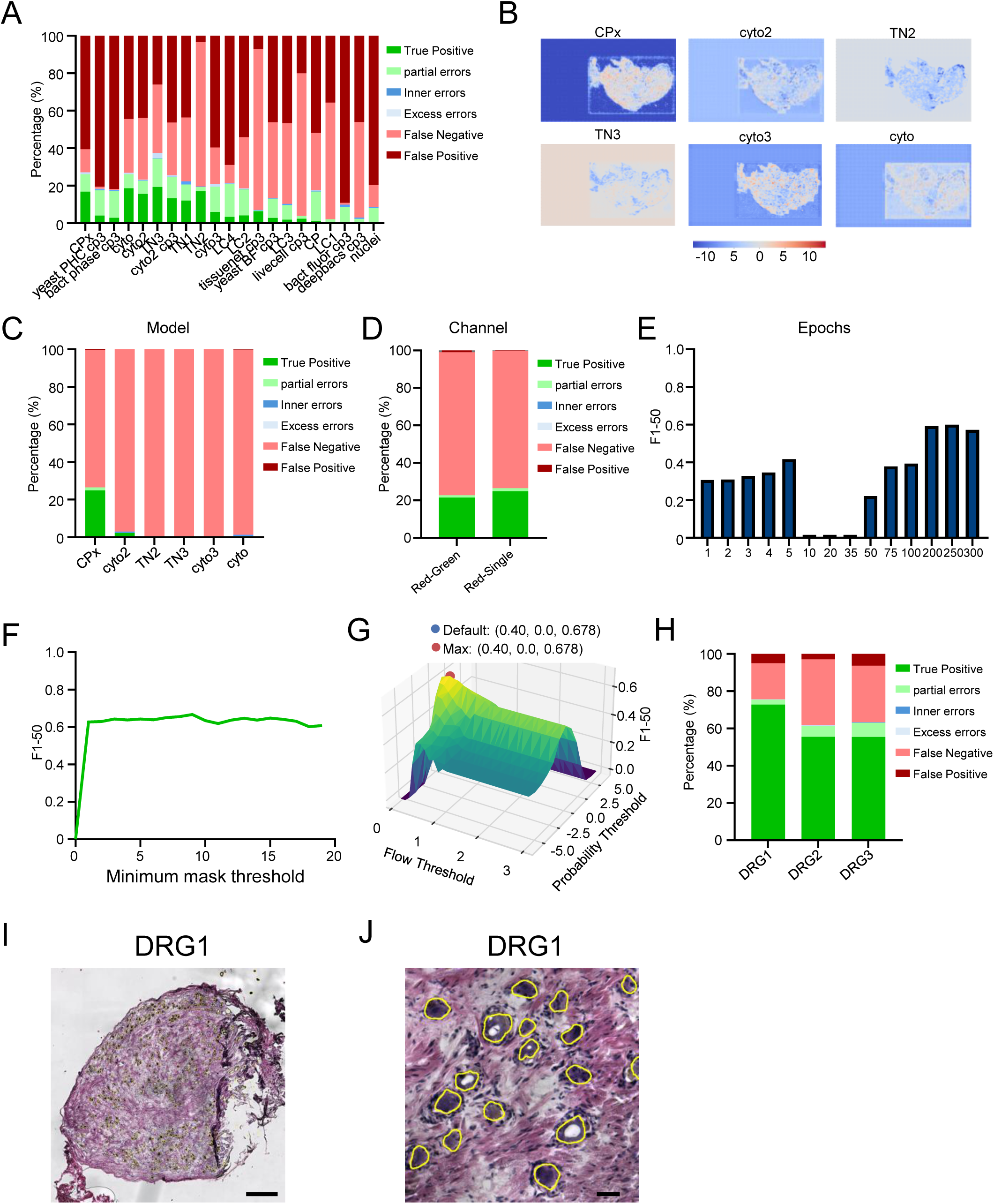
H&E image-based segmentation by Cellpose. (**A**) Segmentation performance evaluation and error classification of all initial pretrained Cellpose models on the DRG4 H&E image without training. (**B**) Visualization of segmentation probability maps from initial top pretrained models (CPx, cyto2, TN2, TN3, cyto3, and cyto) applied to DRG4. (**C**) Segmentation performance on DRG4 of the six top pretrained models after training with DRG1-3 dataset. (**D**) Comparison of segmentation results models with both red and green channels (RG) versus red-only channel (R) using trained CPx. (**E**) Effect of training epochs on segmentation performance for the CPx_R model trained with varying epoch numbers. (**F**) Segmentation performance trained for 300 epochs on CPx_R model across a range of minimum mask thresholds, (**G**) 3D plot of F1-50 scores for CPx _R_9_mask model predictions across a range of flow and probability thresholds, illustrating the optimal parameter combination (Max) compared to the default. (**H**) Segmentation error classification for the Cellpose_H&E model applied to DRG1–3 full tissue images. (**I-J**) Representative segmentation results for DRG1 whole image (I) and a zoomed-in area (J). Predicted neuron boundaries (yellow) overlaid on H&E image. Scale bars: 500 µm in (I) and 50 µm in (J).

**Figure S8.**
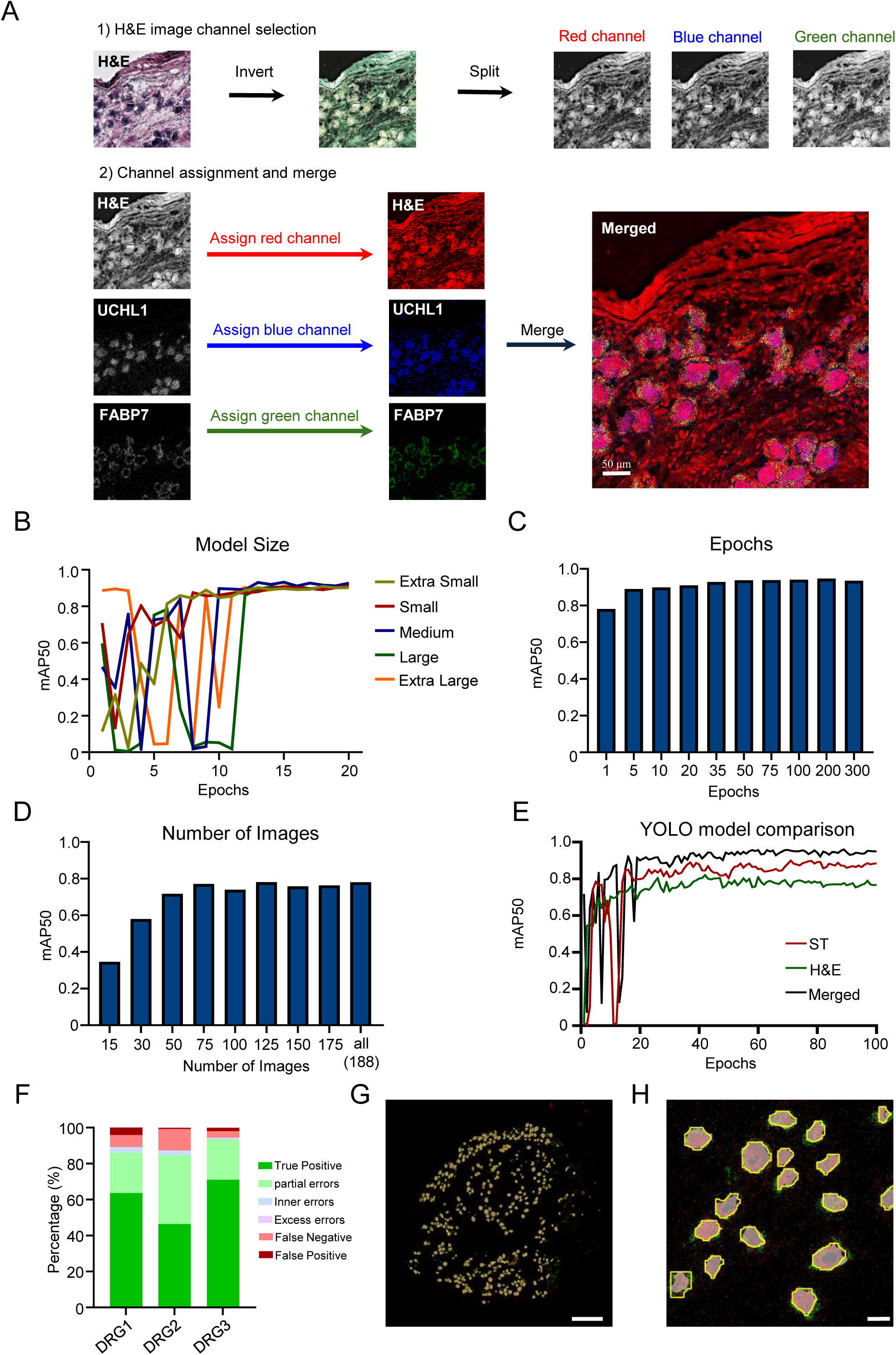
Combine ST and H&E images for segmentation by YOLO. (**A**) Workflow for generating merged images that combine ST and H&E images for YOLO model training. Top: H&E images were inverted and split into red, green, and blue channels. The red channel was selected for merge. Bottom: The selected red channel from the H&E image was combined with ST images—UCHL1 (assigned to blue channel) and FABP7 (assigned to green channel)—to generate a three-channel merged image. Scale bar: 50 µm. (**B**) Effect of model size on segmentation performance. mAP50 values are shown for five YOLO model sizes trained on DRG1–3 merged images over 20 epochs. (**C**) Effect of training epochs on performance. The YOLO small model was trained with 188 images for various numbers of epochs (1 to 300) (**D**) Effect of training set size. The YOLO small model was trained for 20 epochs using increasing numbers of training images. (**E**) Comparison of mAP50 growth during YOLO model training on three image types: ST: transcript-only images, HE: H&E-only images, Merged: combined transcript and H&E images. (**F**) Segmentation error classification for the YOLO_Merged model when applied to DRG1–3 full tissue images. (**G–H**) Representative segmentation results for DRG1 whole image (G) and a zoomed-in area (H). Predicted neuron boundaries (yellow) overlaid on ST image. Scale bars: 500 µm in (G) and 50 µm in (H).

**Figure S9.**
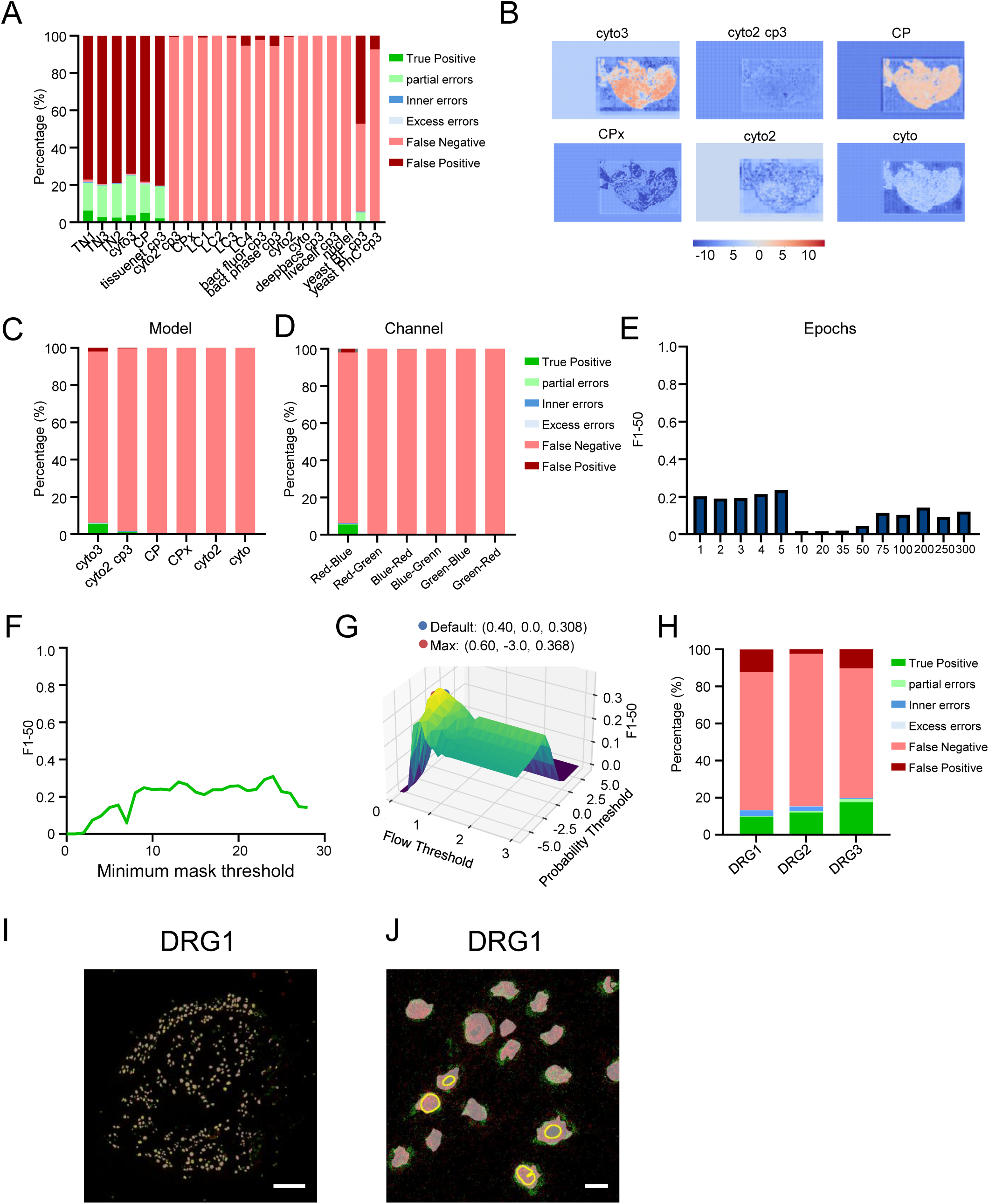
Combine ST and H&E images for segmentation by Cellpose. (**A**) Segmentation performance evaluation of all initial prediction Cellpose models for merged images based on prediction categories applied to DRG4 without training. (**B**) Visualization of segmentation probability maps from initial top pretrained models (cyto3, cyto2 cp3, CP, CPx, cyto2, cyto) applied to merged DRG4. (**C**) Segmentation performance on DRG4 of the six top pretrained models after training with DRG1-3 dataset. (**D**) Comparison of segmentation results trained with different channel combinations using CPx models. (**E**) Effect of training epochs on segmentation performance for the CPx_RB (RB: Red-Blue channel) model trained with varying epoch numbers. (**F**) Segmentation performance trained for 300 epochs on CPx_RB model across a range of minimum mask thresholds, (**G**) 3D plot of F1-50 scores for CPx _ RB_24_mask model predictions across a range of flow and probability thresholds, illustrating the optimal parameter combination (Max) compared to the default. (**H**) Segmentation error classification for the Cellpose_Merged model applied to DRG1–3 full tissue images. (**I-J**) Representative segmentation results for DRG1 whole image (I) and a zoomed-in area (J). Predicted neuron boundaries (yellow) overlaid on ST image. Scale bars: 500 µm in (I) and 50 µm in (J).

**Figure S10.**
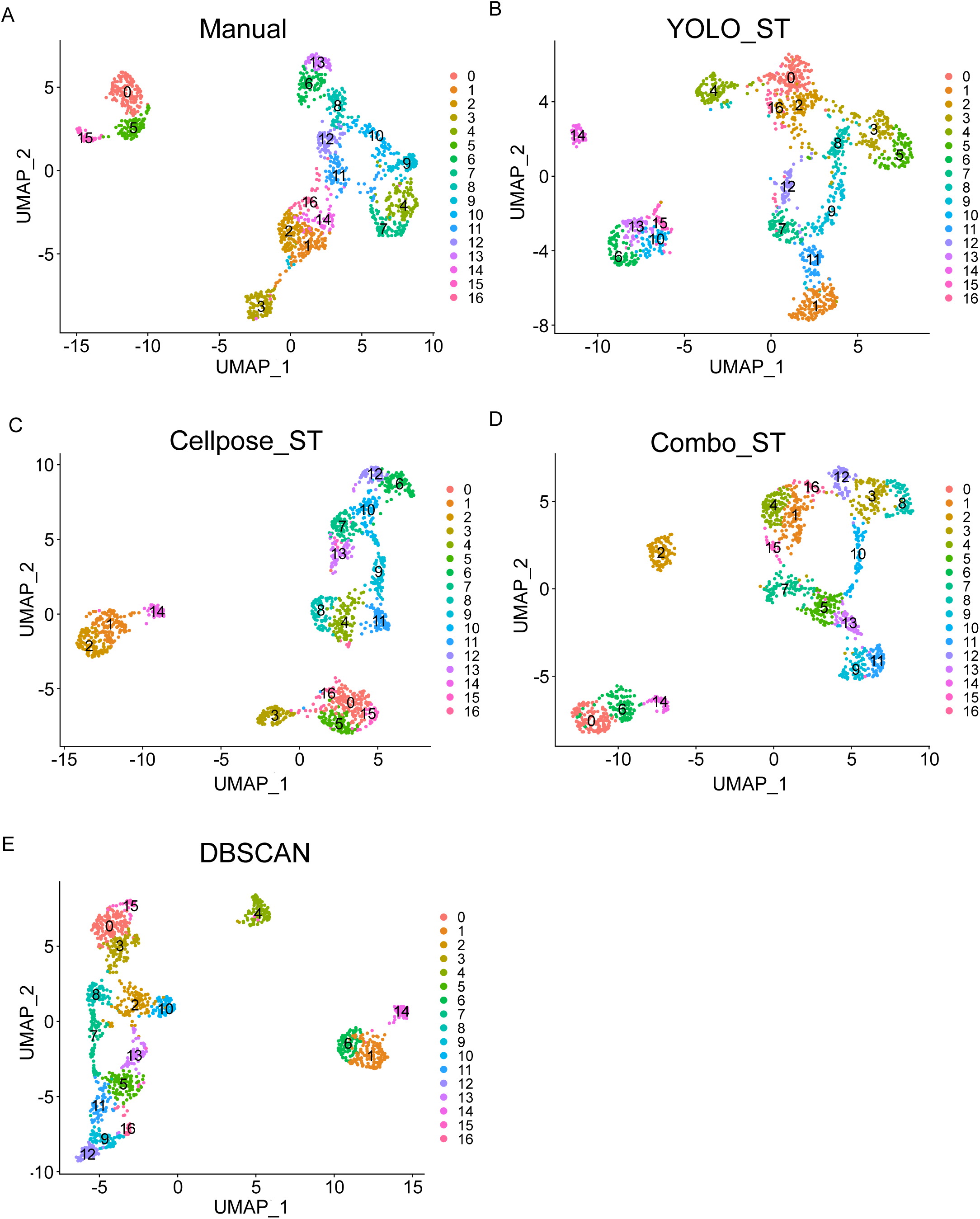
Cell clustering based on segmentation from different models. (**A-E**) UMAP plots showing the clustering of cells from manual annotation (A), Yolo_ST (B), cellpose_ST (C), Combo_ST (D), and DBSCAN (E).

**Figure S11.**
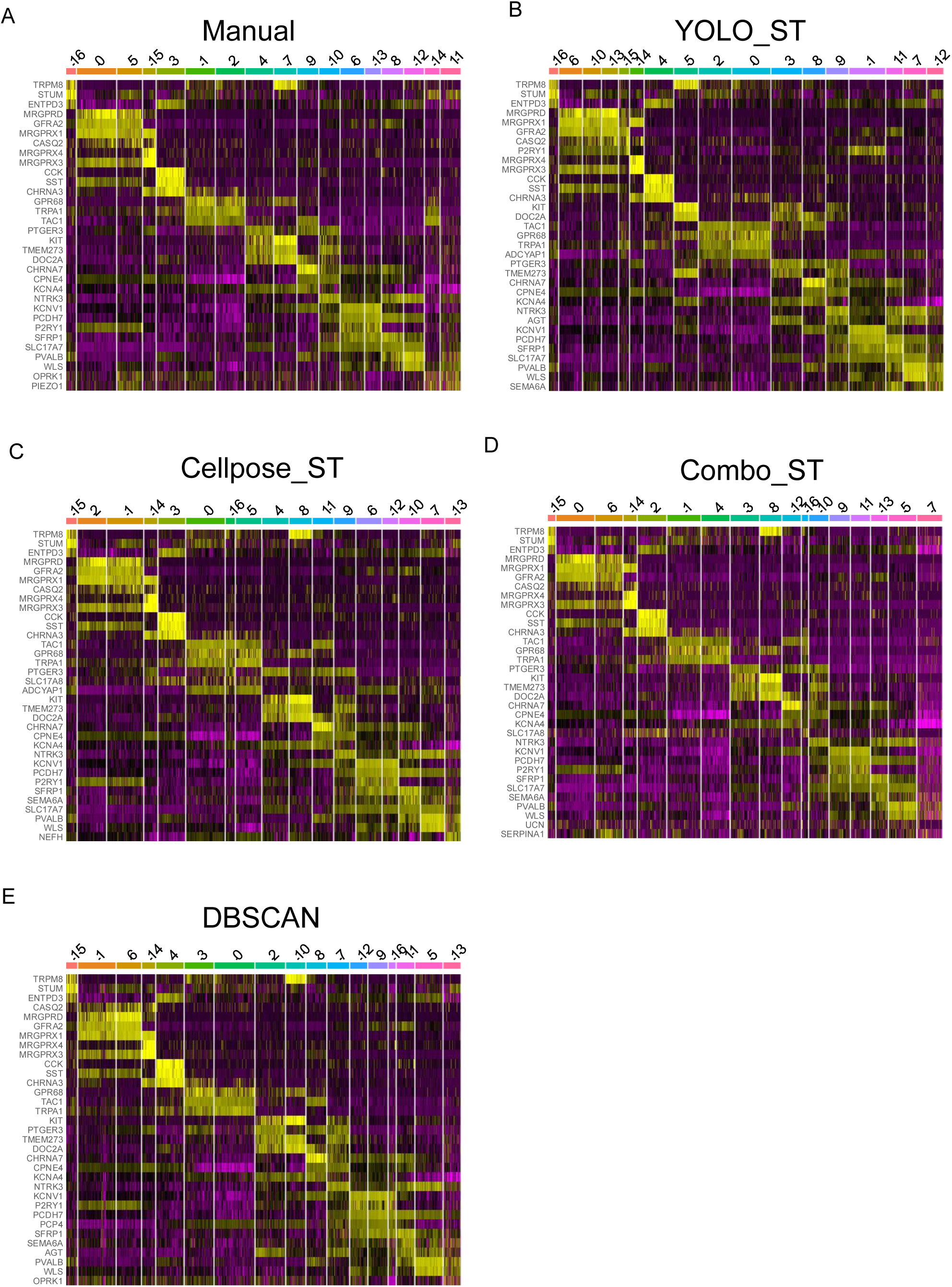
Marker gene identification based on segmentation from different models. (**A-E**) Heatmap plots showing the top marker genes identified for each cluster from manual annotation (A), Yolo_ST (B), Cellpose_ST (C), Combo_ST (D), and DBSCAN (E) segmentation.

**Figure S12.**
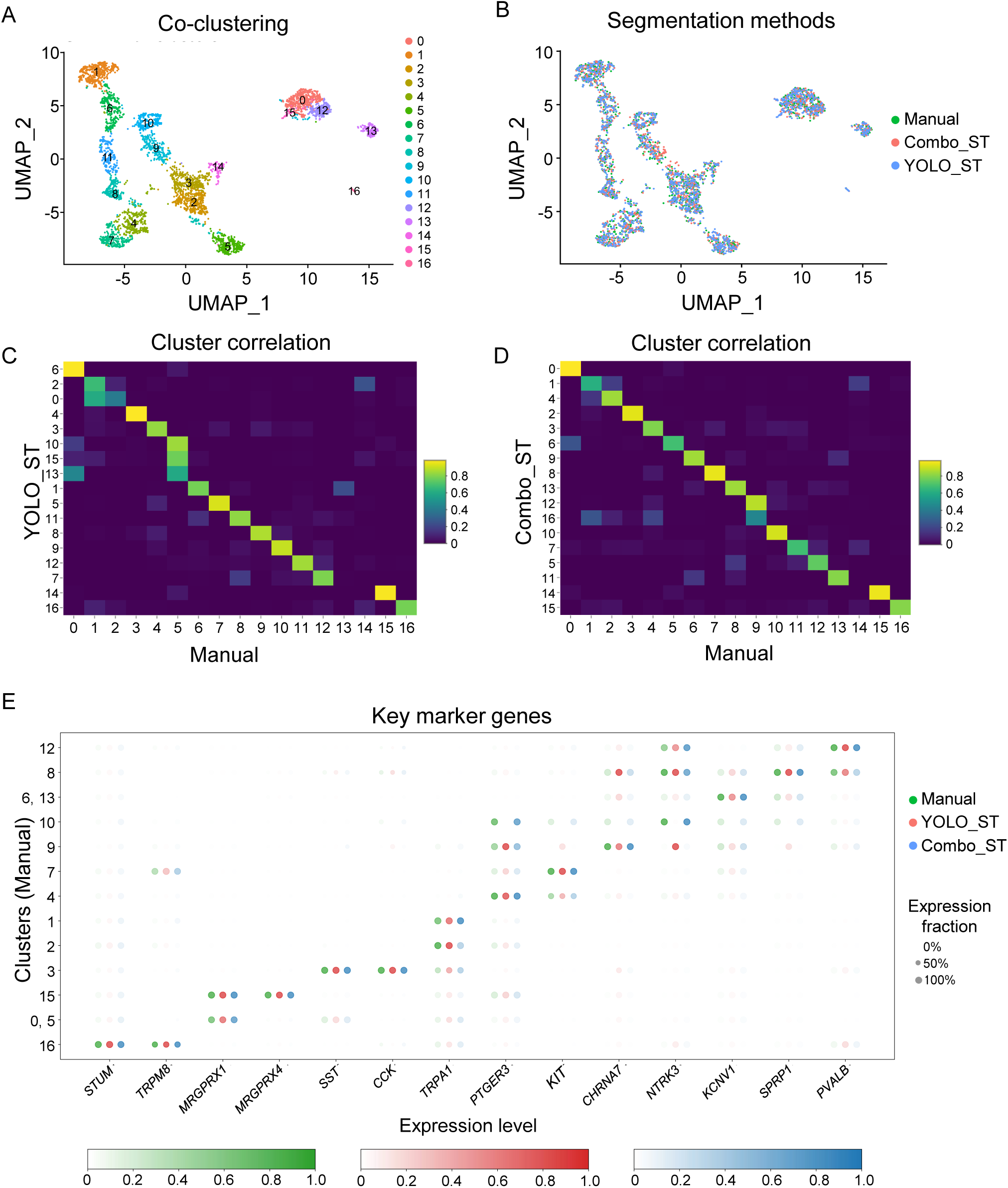
Model comparison with downstream analysis. (**A**) UMAP plot showing the co-clustering of cell segmentation from manual annotation, Combo_ST model, and YOLO_ST model. (**B**) UMAP plot showing the distribution of cell segmentation from manual annotation, Combo_ST model, and Yolo_ST model in each cluster. (**C-D**) Heatmap comparing the similarities of cell clusters between YOLO_ST (C) and Combo_ST (D) segmentation and manual annotation. (**E**) Dot plot showing the top marker genes expressed in each cell cluster from manual annotation (green), YOLO_ST (red), and Combo_ST (blue). The color scale represents the normalized average expression level (from 0 to 1) of each gene in the clusters.

